# Context-Dependent and Gene-Specific Role of Chromatin Architecture Mediated by Histone Modifiers and Loop-extrusion Machinery

**DOI:** 10.1101/2025.02.21.639596

**Authors:** Melodi Tastemel, Adam Jussila, Bharath Saravanan, Hui Huang, Yang Xie, Quan Zhu, Yifan Jiang, Ethan Armand, Bing Ren

## Abstract

Loop-extrusion machinery, comprising the cohesin complex and CCCTC-binding factor CTCF, organizes the interphase chromosomes into topologically associating domains (TADs) and loops, but acute depletion of components of this machinery results in variable transcriptional changes in different cell types, highlighting the complex relationship between chromatin organization and gene regulation. Here, we systematically investigated the role of 3D genome architecture in gene regulation in mouse embryonic stem cells under various perturbation conditions. We found that acute depletion of cohesin or CTCF disrupts the formation of TADs, but affects gene regulation in a gene-specific and context-dependent manner. Furthermore, the loop extrusion machinery was dispensable for transcription from most genes in steady state, consistent with prior results, but became critical for a large number of genes during transition of cellular states. Through a genome-wide CRISPR screen, we uncovered multiple factors that can modulate the role of loop extrusion machinery in gene regulation in a gene-specific manner. Among them were the MORF acetyltransferase complex members (Kat6b, Ing5, Brpf1), which could antagonize the transcriptional insulation mediated by CTCF and cohesin complex at developmental genes. Interestingly, inhibition of Kat6b partially rescues the insulator defects in cells lacking the cohesin loader Nipbl, mutations of which are responsible for the developmental disorder Cornelia de Lange syndrome. Taken together, our findings uncovered interplays between the loop extrusion machinery and histone modifying complex that underscore the context-dependent and gene-specific role of the 3D genome.

## Introduction

Spatial and temporal gene expression is dynamically orchestrated by cis-regulatory elements (cREs) such as enhancers, promoters, insulators, and silencers (*1*). Enhancers control spatiotemporal gene expression from a distance (*2*, *3*). While tremendous progress has been made with the mapping and annotation of candidate enhancers in the genomes of diverse organisms, questions remain regarding how enhancers control specific target genes from a distance. One prominent model posits that enhancers and promoters physically interact through the formation of DNA loops (*4–6*). This DNA looping brings enhancer bound transcriptional activators into proximity with promoter regions and the basal transcription machinery to influence transcriptional dynamics, including the frequency of transcriptional bursting. A number of factors, including CTCF and the cohesin complex have been shown to mediate the formation and maintenance of these loops, in a process known as loop-extrusion (*7*). Additionally, CTCF and the cohesin complex can segregate the genome into chromatin domains known as topologically associating domains (TADs) (*8–12*). It has been shown that loop extrusion could facilitate the enhancer-mediated transcriptional bursting at promoters by stimulating the fusion of RNA polymerase and mediator condensates formed at promoters and enhancers, respectively (*13*). By contrast, TAD boundaries, frequently characterized by convergent CTCF binding sites, constrain interactions between distal enhancers and limit the influence of cis-regulatory elements on gene expression to genes within the same TADs (*1*, *14*). It has been long-held that chromatin organization plays a crucial role in gene regulation. However, this view was challenged by observations that acute depletion of CTCF and cohesin caused gross disruption of local chromatin structure and loss of TADs, but only slight to modest transcriptional changes depending on the cell types studied (*15*). Additional studies have shown that TAD boundaries cannot absolutely prevent an enhancer from interacting or regulating a gene in another TAD (*15*–*17*) and in many cases enhancer strength is also involved in both gene expression levels and sensitivity of a gene to transcriptional insulation (*18*). These observations indicated that the role of chromatin architecture is more complex than previously thought (*19*, *20*), necessitating a comprehensive investigation into how chromatin architecture and insulators operate to modulate selection of target genes by distal enhancers.

To gain insights into the role of chromatin architecture in gene regulation, we conducted systematic investigation of the chromatin organization and gene regulation in mouse embryonic stem cells (mESC). Initially focusing on the Sox2 locus (*21*) whose transcription is driven by a distal enhancer located 110 kb downstream of the gene, we performed single-molecular chromatin tracing experiments and epigenome profiling under various genetic and epigenetic perturbations. We observed a context dependent role of chromatin architecture in the regulation of Sox2 gene expression by its distal enhancers, which highlighted the necessity of enhancer-promoter proximity during establishment of transcriptional states but not maintenance. We then carried out genome-wide analysis of the transcriptional programs and chromatin architecture after acute depletion of cohesin and CTCF under steady state or in differentiation conditions. Some genes behave the way Sox2 did but many more genes show lack of response to cohesin or CTCF depletion. Further, under cell differentiation conditions, a different set of genes showed dependence on CTCF and cohesin. These results suggest gene-specific and context dependent effects of CTCF and cohesin loss. To further identify mechanisms that modulate the context-dependent and gene-specific effects of CTCF and cohesin on enhancer-driven transcription, we conducted a genome wide CRISPR knockout screen and discovered new factors that modulate transcriptional insulation of TAD boundaries. Importantly, we uncovered a role for the MORF histone acetyltransferase complex as an antagonist of transcriptional insulation mediated by CTCF and cohesin loop-extrusion complex. MORF localizes mainly at promoters, where it promotes the formation of promoter-centric DNA loops. Loss of Kat6b subunit of MORF complex resulted in the loss of enhancer-promoter contacts especially at developmental regulator genes, and restoration of transcriptional defects in cells lacking Nipbl, the cohesin loader protein responsible for the Cornelia de Lange Syndrome. Our results therefore suggest the context-dependent and gene-specific role that 3D genome architecture plays in the establishment of new transcription programs during development and pathogenesis, and uncovers novel regulators of this process.

## Results

### Acute depletion of loop-extrusion machinery in steady state mESCs leads to structural changes without affecting transcription at the Sox2 locus

To explore the function of chromatin architecture such as TADs and DNA loops in enhancer dependent gene activation, we utilized a model cell system that we previously reported (*21*). In this system, the two copies of the Sox2 gene were tagged with fluorescent reporters eGFP and mCherry on the CAST and 129 alleles, respectively. An insulator sequence with 4 CTCF binding sites (4xCBS) had been inserted between Sox2 and its super enhancer (110 kb away) on the CAST allele (*21*). This mESC line, referred to hereafter as 4CBS, exhibits reduced eGFP expression and enhancer-promoter contact frequency at the CAST allele as a result of insulator (*21*).

Using 4CBS mESCs, we performed single-molecule chromatin tracing at the Sox2 locus at 5 kb genomic resolution with a multiplexed DNA FISH protocol to identify structural features as described previously (fig. S1a) (*16*, *21*). Our chromatin tracing data revealed high probabilities of TAD boundaries at the Sox2 promoter and super-enhancer (∼10% of single chromosomes) and the 4xCBS insertion site (∼15-20%) (Fig. 1A, *bottom*). Individual chromosomes showed tremendous variability in structure, with Sox2 enhancer-promoter (E-P) distances ranging from the low tens of nm to nearly 1 µm separation (fig. S1b). On average, E-P distances in CAST alleles were greater than those in 129 alleles by ∼35 nm, as a result of the TAD boundary formed at the 4xCBS insertion on this allele.

**Figure 1.**
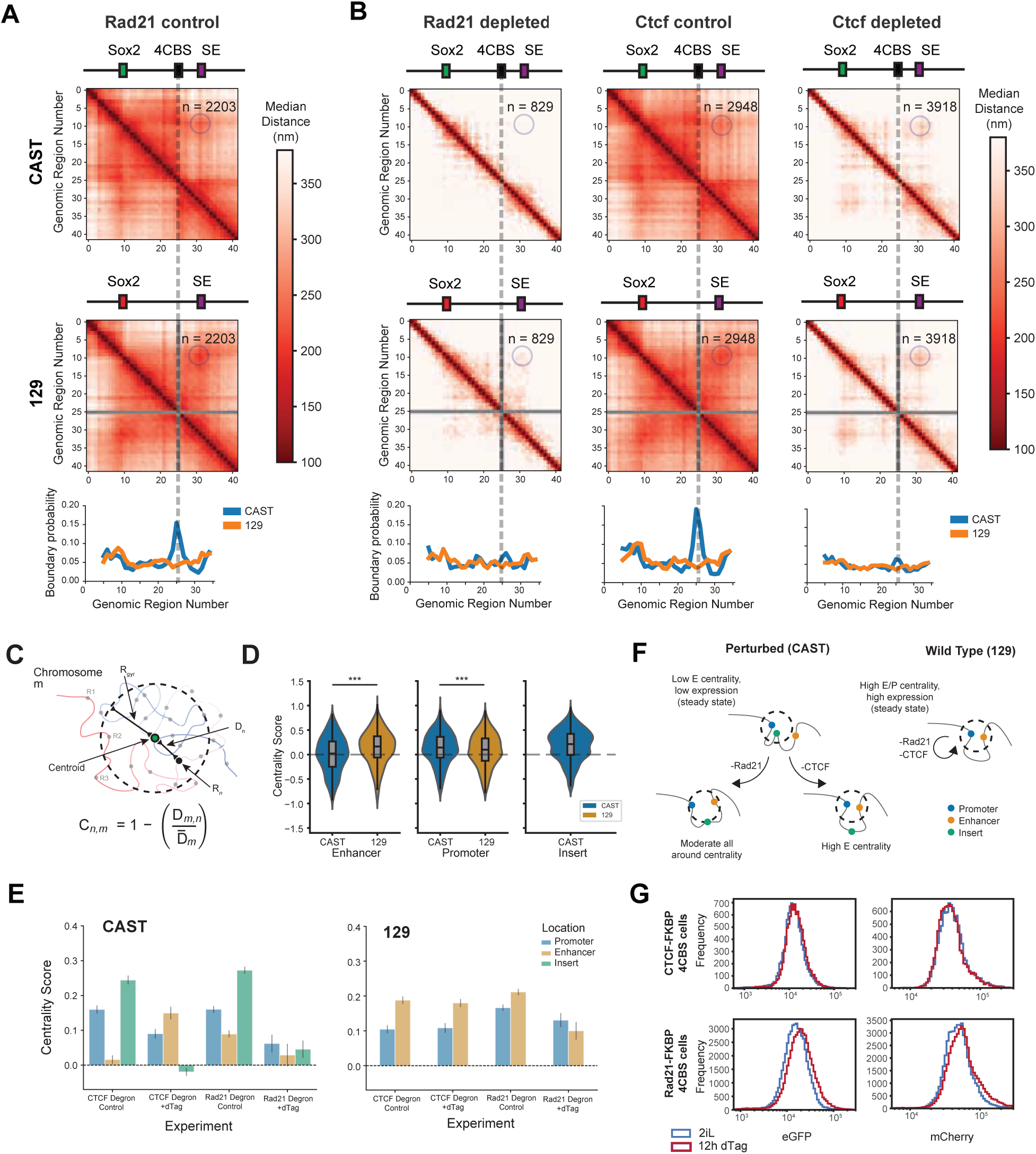
Acute Disruption of TADs in steady state mESCs causes mild transcriptional changes in Steady State mESCs. **(A)** *Upper:* Chromatin tracing median distance matrix for 4CBS Rad21 control cells separated by allele. The dashed line represents the 4xCBS insertion region (absent in the 129 allele), and the circle denotes the location of E-P interaction. *Lower:* Boundary probability computed for each allele from chromatin tracing data. **(B)** *Upper:* Chromatin tracing median distance matrices for Rad21 depleted cells (*left)* CTCF control cells (*middle)*, and CTCF depleted cells (*right)*. The dashed lines represent the 4xCBS insertion region, and the circle denotes the location of E-P interaction. *Lower:* Boundary probability computed for each allele from chromatin tracing data. (**C)** Cartoon depicting the method of computing centrality score, Cn,m. (**D)** Violin plots of centrality scores from Rad21 degron control chromatin tracing showing distributions of enhancer, promoter, and insert regions. Enhancer centrality is more enriched in the 129 allele than the CAST (p-value = 1.6e-194). Promoter centrality enrichment is present for both alleles, though it shows higher enrichment in the 129 allele (p-value 2.3e-131). Insertion is enriched for centrality in the CAST allele. (**E)** Bar plot showing median centrality scores (derived from single chromosome centrality score distributions as shown in D) per condition for enhancer, promoter, and insert regions on each allele. (**F)** Cartoon depicting conformational change upon CTCF or Rad21 depletion in CAST allele with minimal changes in conformation of 129 allele. (**G)** Histograms of flow cytometry data for control and 12 hour depletion conditions for Rad21 and CTCF degron cells lines. Values indicate normalized expression of Sox2 RNA (eGFP and mCherry).

To determine how disruption of TADs would affect the chromatin structure, enhancer-dependent transcription of Sox2 as well as transcriptional insulation at the insulator, we performed acute depletions of CTCF and cohesin complex subunit Rad21 using independently derived 4CBS mESC lines where Rad21 or CTCF gene has been genetically engineered to carry a FKBP12^F36V^ degron sequence in both alleles (fig. S2a, b). Chromatin tracing analysis upon depletion of either CTCF or Rad21 (Fig. 1A, B) revealed significant loss of long range contacts in both Sox2 alleles, dramatically reduced boundary probability at the 4xCBS insertion, consistent with the role of CTCF and cohesin in the formation of TADs (*22*, *23*). E-P distances increased by approximately 100 nm and 150 nm upon CTCF and Rad21 depletion respectively (fig. S2c, d). Despite this, we still observed slight enrichment for E-P contact relative to other regions in the Sox2 locus (Fig. 1A, B). In addition, the allelic median E-P distance differences between CAST and 129 alleles (∼30 nm) were maintained upon CTCF/Rad21 depletions despite the loss of the TAD boundary (Fig. 1A, B). To examine the effect of acute CTCF/Rad21 depletion on Sox2 expression, we performed flow cytometry. We observed no significant changes in Sox2 expression from either allele upon CTCF depletion (Fig. 1G, *top row*). Interestingly, Rad21 depletion resulted in a subtle increase in Sox2 expression from both CAST and 129 alleles (Fig. 1G, *bottom row*), which we attributed to increased mixing of accessible chromatin (*24*, *25*). RNA-seq also revealed minimal changes in expression for Sox2 and genome-wide upon either short-term acute Rad21 or CTCF depletions, consistent with previous literature (*15*, *22*, *26*) (fig. S2f). These results were consistent with previous reports that cohesin and CTCF loss caused slight transcriptional changes in mESC and other cell types (*15*, *22*, *23*, *26*).

To better understand the relationships between chromatin structure and gene expression, we looked into the heterogeneity of chromatin structure after perturbations, by applying dimensionality reduction methods using the pairwise distances on each chromosome as features (for details see Methods). We found that chromosome structures were in a highly dynamic continuum with only mild allelic biases (fig. S1c) (*19*, *27*), highlighting the need for alternative analyses to understand single chromosome structural changes. For this reason, we chose to quantify structures by their position of regions relative to the centroid of each chromosome instead of euclidean distances alone (Fig. 1C). Earlier studies utilized centrality to highlight regions extruded by cohesin or bound by CTCF, showing how a region with high centrality within a domain naturally leads to increased contacts with all other regions in the same domain (*19*, *20*). Centrality score distributions for the enhancer, promoter, and insertion were distinct between alleles (Fig. 1D; fig. S1d). In the 129 allele, we identified centrality enrichment for both the promoter and enhancer regions (∼10% and 20% enrichment respectively). On the other hand, CAST alleles displayed a notable centrality enrichment for the insertion and the promoter regions (∼22% and ∼15%). Example chromosome traces displaying both high and low centrality show what such structures might look like (fig. S1e).

Following CTCF depletion, centrality score measurements revealed a distinct conformational switch in the CAST allele, where the centrality of enhancer and 4xCBS insert were swapped (Fig. 1E). The resulting high enhancer and promoter centrality conformation was similar to that of the control 129 allele (Fig. 1A) and 4CBS mutant (4xCBS removed) (fig. S2e) (*21*). Rad21 depletion, on the other hand, resulted in a general loss of centrality locus-wide. This suggests that cohesin looping could play a more general role in mediating centrality than CTCF. In contrast to the insulated CAST allele, the co-centrality of the enhancer and promoter for the non-insulated 129 allele seemed to be independent of both CTCF and Rad21 (Fig. 1E, *right*; Fig. 1F).

### CTCF and Cohesin are required for transcriptional insulation during state transition

The above findings showed that depletion of CTCF and cohesin complex can disrupt the TADs and enhancer-promoter loop at the Sox2 locus without affecting transcriptional insulation, but numerous studies have highlighted the importance of TADs and chromatin structure in development and gene expression (*28–33*). We hypothesized that the TADs and loops could become essential for setting up the transcriptional state of a gene when it is reprogrammed or reset.

To test this possibility, we shifted the 4CBS mESCs from serum free 2i media known for maintaining naive pluripotentcy to serum containing media without LIF (*34*, *35*) for 24hrs to induce onset of differentiation (Fig. 2A). Flow cytometry showed that the insulated Sox2-Egfp allele was suppressed compared to the Sox2-mCherry allele due to transcriptional insulation by the 4xCBS insulator. Contrary to the results in steady state, Sox2-eGFP allele is robustly activated after CTCF depletion during differentiation (Fig. 2B). Chromatin tracing revealed that, consistent with steady state, CTCF depletion in differentiation media also resulted in a loss of TAD boundary at the 4xCBS insertion (Fig. 2C) and resulted in a centrality switch (Fig. 2D, *left*). Unlike in steady state, the locus retained contacts in differentiation media, including interactions between the Sox2 gene and the super-enhancer (Fig. 2C) and did not increase the E-P distances upon CTCF loss (fig. S3a). Correlating E-P distances, promoter centrality, and enhancer centrality across perturbations and alleles with RNA expression, we found that enhancer centrality showed the highest correlation with RNA-seq data (p=0.83) (Fig. 2E). Consistent with the transcriptional readout, H3K27ac at the Sox2 promoter increased at the Sox2 promoter in the insulated allele upon CTCF depletion after differentiation, but not in the steady state (Fig. 2F). We also acutely depleted cohesin subunit Rad21 and observed similar reactivation of Sox2-eGFP allele along with increased H3K27ac, albeit slightly, at the Sox2 promoter (Fig. 2G; fig. S3b). These observations support our hypothesis that loop-extrusion machinery and chromatin organization plays a role in gene regulation and transcriptional insulation during transition of cellular state.

**Figure 2.**
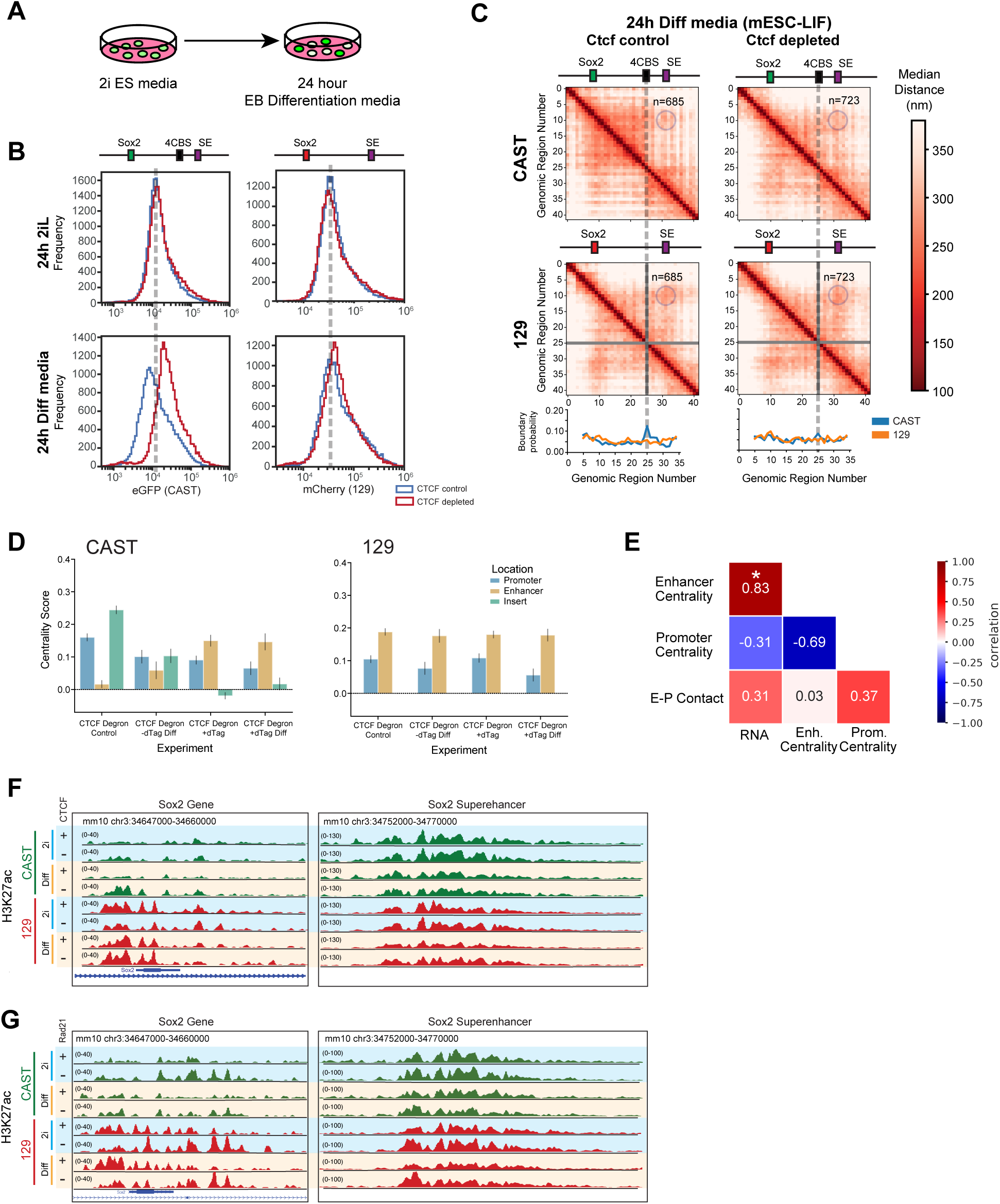
Onset of differentiation demonstrates the need for loop extrusion to achieve new transcriptional states. **(A)** Schematic of induction of differentiation for 4CBS cells. Cells grown in 2i embryonic stem cell media and transferred to embryoid body differentiation media for 24 hours. **(B)** Histograms of flow cytometry data for control and CTCF depletion conditions in 2i and differentiation media showing normalized Sox2-eGFP and mCherry expression from CAST and 129 allele respectively. The dashed lines indicate the approximate location of the peak in control 4CBS cells for visual reference. (**C)** *Upper:* Median distance matrices for CAST and 129 alleles in CTCF control and CTCF depletion for cells grown in differentiation media for 24h. *Lower:* Boundary probability computed for each allele from chromatin tracing data. **(D)** Barplot showing median centrality scores for various conditions for enhancer, promoter, and insert regions on each allele. **(E)** Correlation heatmap comparing normalized RNA-seq counts with enhancer centrality, promoter centrality, and E-P contact across alleles and perturbations (includes WT and differentiation media). Boxes are colored and labeled by pearson correlation coefficients (−1 to 1) and marked for significance. **(F)** Genome browser tracks showing the H3K27ac ChIP-seq signal at the promoter and enhancer in each condition (2i or differentiation media and +/− dTag) in CTCF degron 4CBS cells for both the CAST and 129 alleles. **(G)** Genome browser tracks showing the H3K27ac ChIP-seq signal at the promoter and enhancer in each of the conditions (2i or differentiation media and +/− dTag) in Rad21 degron 4CBS cells for both the CAST and 129 alleles.

To further evaluate whether loop-extrusion machinery is necessary for transcriptional regulation by TAD boundaries during transition of cellular states, we artificially induced a transcriptional state reset by inhibiting histone acetyltransferase p300/CBP, which is required for enhancer dependent transcriptional activation in mammalian cells, using the p300/CBP inhibitor A485 (*36*). We then removed the inhibitor to allow the transcriptional state to re-establish (Fig. 3A). Upon inhibition of p300 activity, both CAST and 129 alleles showed a significant loss of Sox2 expression (fig. S4a). Upon removal of the inhibitor, we observed that the 129 allele recovered to near-previous expression levels after 12 hours (Fig. 3B; fig. S4a, b). In contrast, the CAST allele did not show the same recovery behavior, with expression staying well below normal CAST levels, likely due to the transcriptional insulation mediated by the 4xCBS insulator. Indeed, acute depletion of CTCF reactivated the expression of Sox2-eGFP from the CAST allele (Fig. 3B). In accordance, H3K27ac levels were elevated at the CAST allele promoter early in the p300 activity recovery with CTCF depletion, but not in its presence (Fig. 3C).

**Figure 3.**
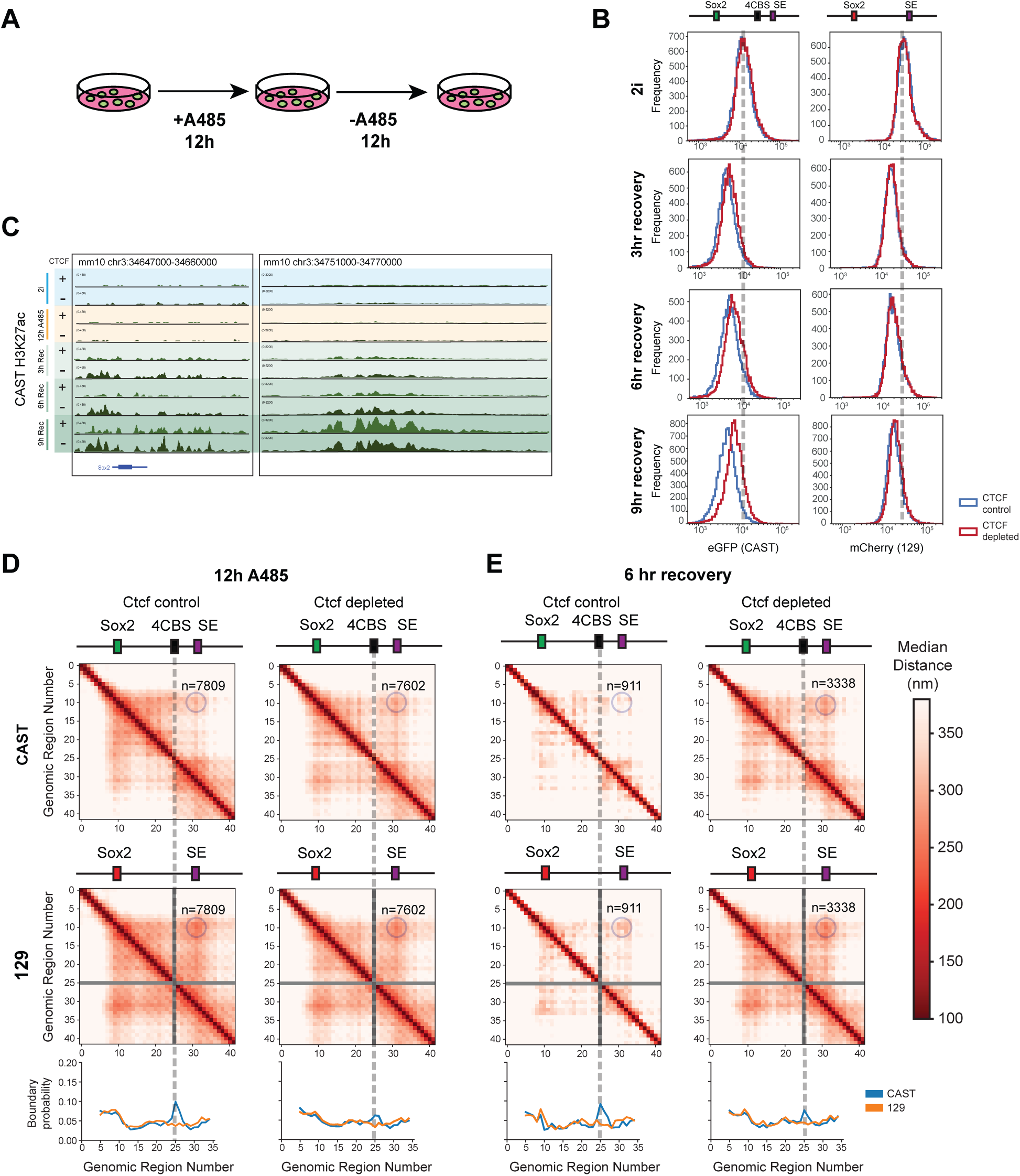
TADs are required for transcriptional insulation during recovery from acute epigenetic perturbation. **(A)** Cartoon depicting the A485 treatment and recovery. **(B)** Histograms of flow cytometry data for control and CTCF depletion conditions in 2i as well as 3, 6, and 9 hours of A485 recovery showing normalized Sox2-eGFP and mCherry expression from CAST and 129 allele respectively. The dashed lines indicate the approximate location of the peak in control 4CBS cells for visual reference. **(C)** Genome browser tracks showing H3K27ac ChIP-seq signal at the promoter and enhancer for eGFP-CAST allele in control 2i media, 12hr A485 treatment, and 3, 6, and 9 hours A485 recovery following 12hr A485 treatment. **(D)** *Upper:* Median distance matrices for CAST and 129 alleles after 12 hours of A485 treatment in both CTCF control and CTCF depletion. *Lower:* Boundary probability computed for each allele from chromatin tracing data. **(E)** *Upper:* Median distance matrices for CAST and 129 alleles after 12 hours of A485 treatment and 6 hours of recovery (removal of A485) in both CTCF control and depleted conditions. *Lower:* Boundary probability computed for each allele from chromatin tracing data.

We further assessed structural changes at the Sox2 locus using chromatin tracing. p300/CBP inhibition did not result in significant structural changes in either Sox2 allele upon inhibition of p300/CBP. By contrast, CTCF depletion during inhibition and recovery resulted in the weakening of the 4CBS boundary strength and a conformation of enriched enhancer and promoter centrality (Fig. 3D, E; fig. S4c). Loss of CTCF during the recovery stage resulted in greater enhancer E-P contacts (Fig. 3E, *right*). We confirmed that the recovery of Sox2 expression was dependent on the Sox2 Super Enhancer (SE) using allelic SE deletions (fig. S4d, e). Similarly, cohesin depletion during the same state reset as above led to re-expression of insulated Sox2 on the CAST allele (fig. S4f).

Collectively, these results demonstrated that although disruption of TADs resulted in no transcriptional changes at the Sox2 locus in steady state, it led to loss of transcriptional insulation during cell state transitions such as onset of differentiation or recovery from P300/ CBP inhibition. The results support a model in which TADs and transcriptional insulators are required to direct specific enhancer-promoter communications during establishment of a new transcriptional state but could be dispensable in steady state.

### Gene-specific role of loop extrusion machinery during cellular state transitions

To better understand the global impact of disruption of TADs and chromatin loops on gene expression, we performed bulk RNA-seq and measured fold changes in expression for each gene upon CTCF or Rad21 depletion in steady state or during cell differentiation (Fig. 4A-E). Overall, we found that 1970 genes were significantly down-regulated (p<0.05), while 1775 were significantly induced upon CTCF depletion. This was comparable to 24h CTCF loss at steady state (Fig. 4A, B). When comparing the differentially expressed genes (DEGs) upon CTCF depletion in 2i and differentiation media, we found that 49% of genes (1813/3716) showed expression changes in the same direction upon depletion in either context (expression up in both or down in both 2i and differentiation media upon CTCF depletion) (Fig. 4C).

**Figure 4.**
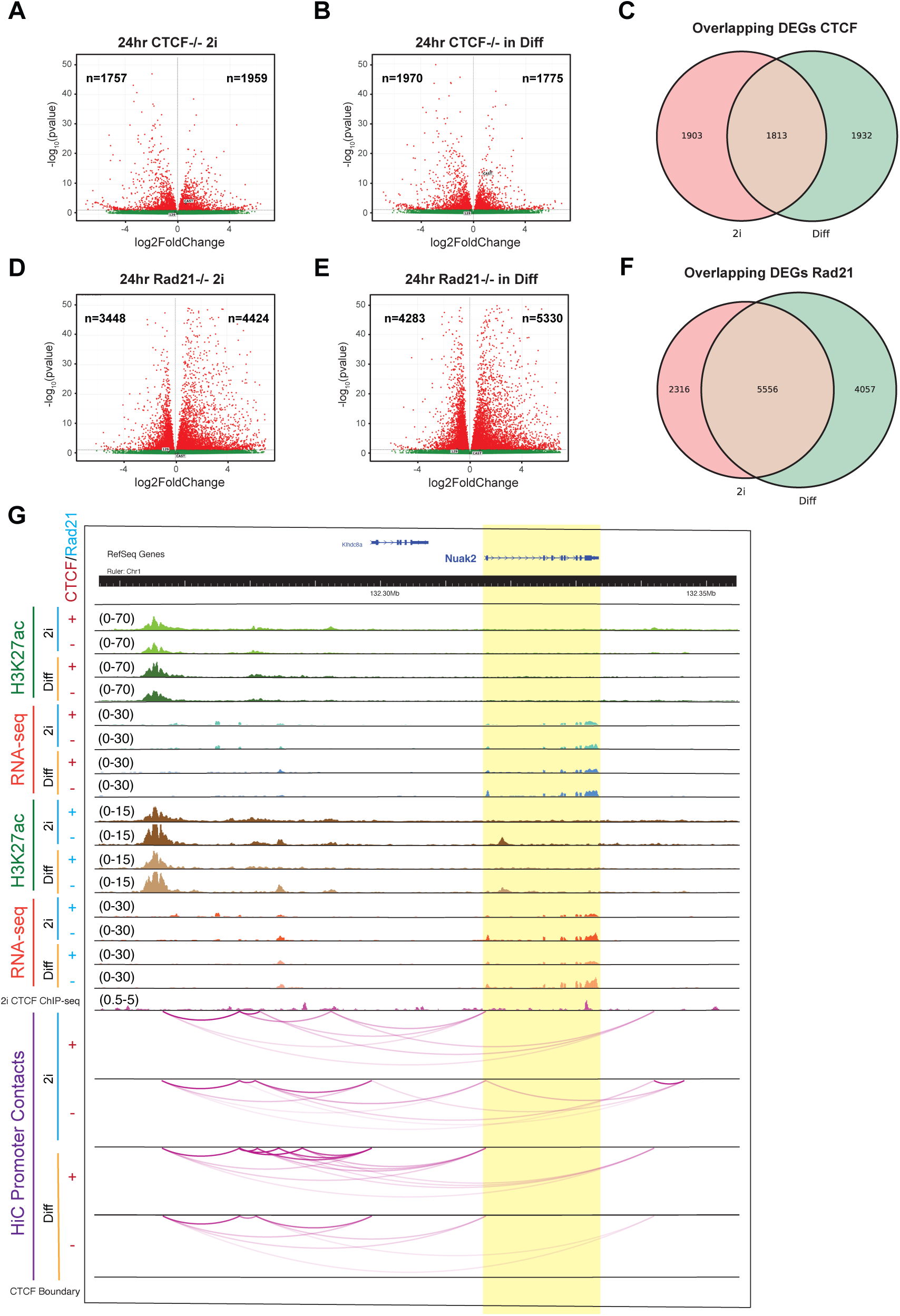
Genome-wide transcriptomic analysis reveals a subset of Sox2-like genes among other diverse 3D genome contexts. **(A)** Volcano plots showing the number of significantly changing genes from bulk RNA-seq data upon 24 hr CTCF depletion in 2i media conditions. Red dots indicate a significance of p-value < 0.05. **(B)** Volcano plots showing the number of significantly changing genes from bulk RNA-seq data upon 24 hr CTCF depletion in differentiation media conditions. Red dots indicate a significance of p-value < 0.05. **(C)** Venn diagram showing the proportion of DEGs which change in the same direction across 2i and Diff for CTCF degron cells. **(D)** Volcano plots showing the number of significantly changing genes from bulk RNA-seq data upon 24 hr Rad21 depletion in 2i media conditions. Red dots indicate a significance of p-value < 0.05. **(E)** Volcano plots showing the number of significantly changing genes from bulk RNA-seq data upon 24 hr Rad21 depletion in differentiation media conditions. Red dots indicate a significance of p-value < 0.05. **(F)** Venn diagram showing the proportion of DEGs which change in the same direction across 2i and Diff for Rad21 degron cells. **(G)** Genome browser showing changing H3K27ac, expression, and contact at the gene Nuak2 across differentiation and removal of CTCF, or Rad21 matching patterns of change seen in the Sox2 CAST allele.

In contrast, 24h Rad21 depletion combined with onset of differentiation resulted in 4283 genes showing a significant negative fold change while 5330 showed a positive fold change, a substantial increase over 24h Rad21 loss at steady state (Fig. 4D, E). When comparing the DEGs upon Rad21 depletion in 2i and differentiation media, we found that 71% of genes (5556/7872) changed expression in the same manner in both 2i and differentiation media (Fig. 4F). Combined with increasing numbers of DEGs, this implied that the magnitude of the changes was likely greater in the transition state than in steady state for Rad21 depletion. Due to our observations of substantially different effects in CTCF and Rad21 depletion conditions, we wanted to understand the extent to which individual genes showed agreement between perturbations. We found that the number of genes with the same direction of expression changes in both CTCF and Rad21 depletion had higher overlap in the genes with induced expression, highlighting a likely link to ectopic activation of genes by previously inaccessible enhancers (fig. S5a, b).

Many genes with unique expression and acetylation changes were identified across our perturbations with dependencies on CTCF and/or Rad21. One example, Nuak2, is located ∼50 kb downstream of a SE and it experienced mild increases in gene activity upon removal of CTCF in 2i and significant increases in differentiation media (Fig. 4G). Additionally, upon depletion of Rad21 in either 2i or differentiation media we see a large increase in expression which is not dependent on the state transition. Much like our Sox2 CAST allele, Nuak2 gene shows a dependence on CTCF to block a nearby SE in WT cells. Additionally, there are genes whose differential expression is affected predominantly by loss of either CTCF alone (fig. S5c) or Rad21 alone (fig. S5d).

While we observed a clear context dependence for the expression of Sox2 in the insulated CAST allele, our RNA-seq analysis showed 691 genes behaved similarly to the Sox2-eGFP allele (differentially expressed alone only in differentiation media without CTCF). These genes do not share any other obvious connection to each other. This seems to further imply a gene-specific dependence for insulation effects, where the role of insulators (enhancing or inhibiting expression) depends on both the gene in question and the cellular context (differentiation stage, enhancer association, insulator position, cell type, etc). It also raises the question of which factors then regulate insulator binding and application in each gene context to impart specificity to each gene promoter or E-P pair.

### Genome-wide CRISPR screen identified regulators of transcriptional insulation

The observation of context-dependent and gene-specific effects of chromatin architecture led us to hypothesize that there are likely modulators that determine how loop extrusion machinery and TAD boundaries affect transcriptional insulation and enhancer-dependent gene activation in different contexts (*37*). Therefore, we carried out a genome wide CRISPR screen using the 4CBS mESC line to identify loss-of-function mutants that either enhanced or dampened transcriptional insulation of the insulated Sox2-eGFP allele (Fig. 5A). Two independent 4CBS clones were transduced with pooled lentiviral sgRNA library (*38*) at a low MOI (∼0.3) so that each infected cell expressed a single sgRNA. The cells were then selected, subsequently expanded, and sorted into two populations: Sox2-eGFP-dim (bottom 10%) and Sox2-eGFP-high (top 10%). The relative sgRNA representations in each population were determined using next generation DNA sequencing (see methods). By comparing the Sox2-eGFP-dim and Sox2-eGFP-bright populations to the control cells, we were able to identify factors that (1) enhance or (2) diminish CTCF insulator function (Fig. 5B, C) (*39*). From these, we filtered the top hits for GFP-bright and -dim populations using a p-value cutoff of 0.05.

**Figure 5.**
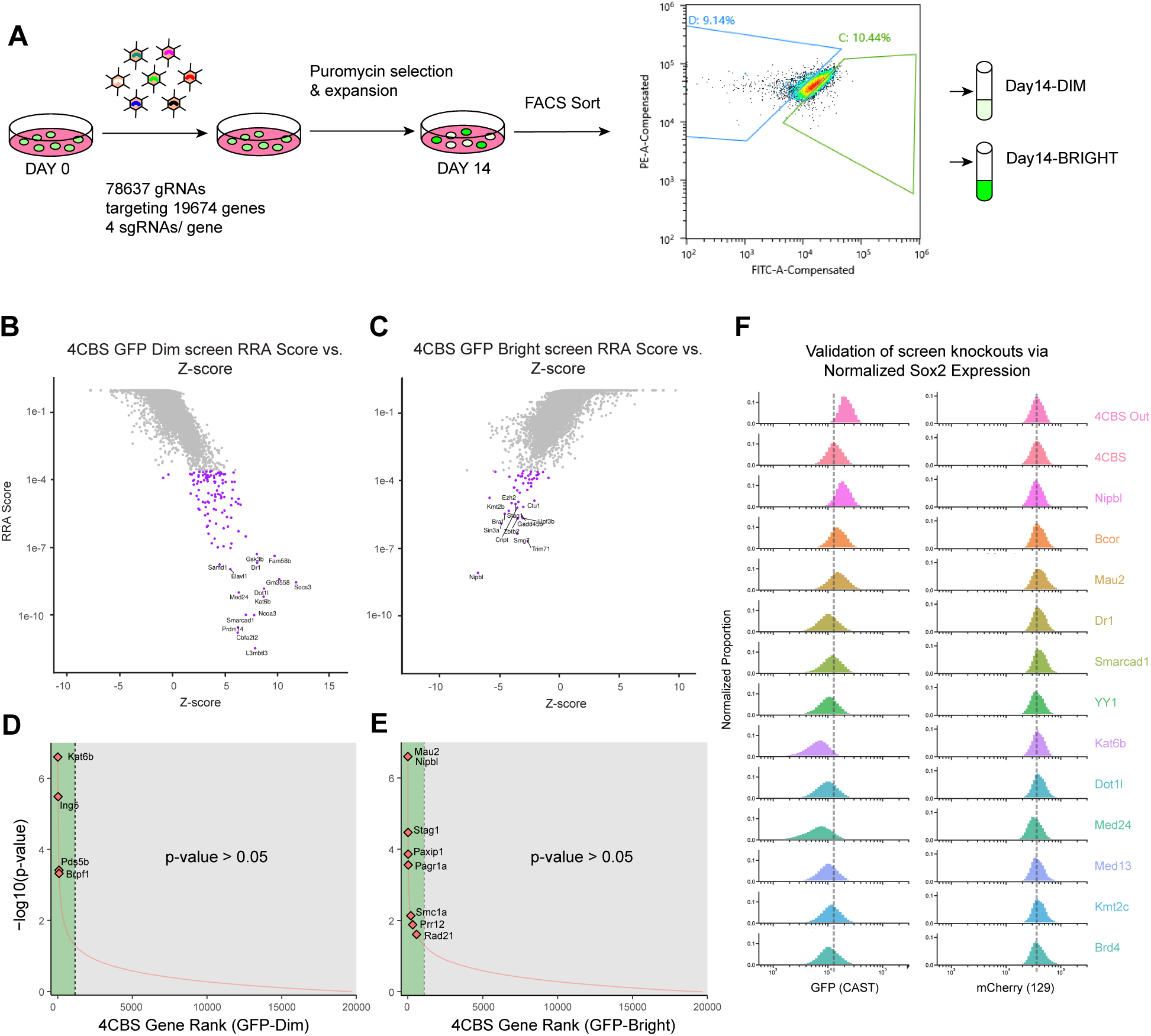
Genome-wide CRISPR Screen Identified Transcriptional Insulation Regulators. **(A)** Schematic depicting GECKO screen scheme. **(B, C)** Scatterplot of MAGeCK screen analysis RRA scores versus z-scores for GFP-DIM (B) and GFP-BRIGHT (C) populations, with FDR < 0.05 genes highlighted in purple. Select genes randomly labeled among the significant genes. **(D, E)** Rank plot showing gene rank vs. p-value for 4CBS cells, highlighting select genes. Green indicates p-value < 0.05 significance threshold, gray represents non-significant hits. (D) GFP-DIM population, MORF acetyltransferase complex subunits highlighted among top hits. (E) GFP-BRIGHT population, cohesin complex and associated proteins highlighted among top hits. (**F)** Histograms of normalized eGFP or mCherry expression from individual lentiviral CRISPR knockout of select significant 4CBS screen hits. The dashed lines aligned with parental 4CBS cell expression of either Sox2 eGFP or mCherry.

Several lines of evidence support the biological relevance of the factors identified in this screen. First, we repeated the screen for Sox2 dual tagged mESCs that lacked the insulator insertion on the CAST allele, and found that most of the top hits identified (fig. S6a, b), except for general transcriptional regulators such as mediator subunits (Med24, Med13, etc) and RNA Polymerase subunits (Polr2g, Polr3c, etc.) (fig. S6c, d) were no longer significant. Second, many cohesin complex subunits, namely Rad21, Stag1, Nipbl, and Mau2 were identified as significant hits (Fig. 5D, E). We also recovered proteins known to interact with cohesin members and have roles related to cohesin function and chromatin occupancy, such as Paxip1, Pagr1a and Prr12 (*40–43*). Additionally, we were able to recover Pds5b, which interacts with Wapl to mediate cohesin release from chromatin (*44–47*) in our GFP-low fraction. Third, we also confirmed the top hits using genetically engineered mESC cells with loss-of-function of individual genes (Fig. 5F).

### Kat6b / MORF complex antagonizes transcriptional insulation by regulating loop extrusion machinery

The finding of 3 out of 4 members of the MORF histone acetyltransferase complex (Kat6b, Ing5 and Brpf1) among the top hits in the GFP-dim group suggested a role of this histone acetyltransferase complex in transcriptional insulation by the loop extrusion machinery (Fig. 5D). We hypothesized that Kat6b (MORF) could antagonize transcriptional insulation mediated by loop extrusion machinery and TAD boundaries. Kat6b, as the enzymatic subunit of MORF, is a member of the MYST family of acetyltransferases and catalyzes H3K14/23 acetylation (*48*). siRNA knockdown experiments on 4CBS and 4CBS insulation mutant mESCs (Fig. 6A,B) confirmed that Kat6b knockdown further enhanced Sox2-eGFP insulation and that this effect depended on having a functional insulator. Similarly, targeting the Ing5 subunit with lentiviral sgRNAs also led to enhanced transcriptional insulation (fig. S7a). To further determine the effect of Kat6b loss on Sox2 locus chromatin structure, we generated Kat6b knockout mESCs (fig. S7b) and performed chromatin tracing (fig. S7d, e). Consistent with the observed enhanced transcriptional changes, Kat6b loss weakened the enhancer-promoter contacts on the CAST allele (fig. S6b, *bottom*) without significant effect on 4CBS boundary formation (fig. S7d, e).

**Figure 6.**
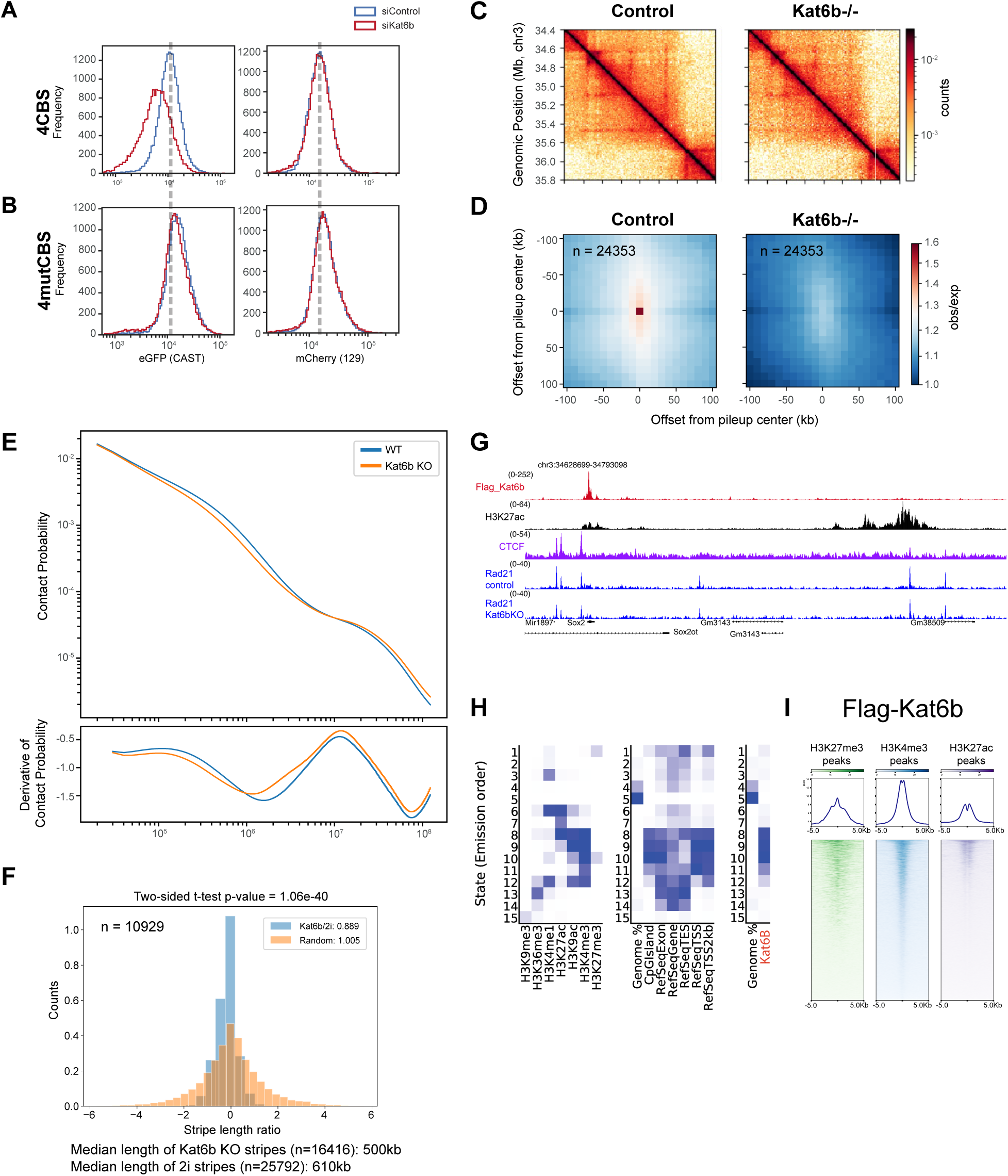
Kat6b / MORF Regulates Transcriptional Insulation by Affecting Loop Extrusion. **(A)** Flow cytometry data showing normalized GFP expression for Kat6b siRNA knockout and Kat6b siRNA control experiments in 4CBS cells. The dashed line represents the approximate median value of the siRNA control for visual aid. (**B)** Flow cytometry data showing normalized GFP expression for Kat6b siRNA knockout and Kat6b siRNA control experiments in 4CBS mutant cells. The dashed line represents the approximate median value of the siRNA control for visual aid. (**C)** Hi-C contact matrix showing an example locus where Kat6b KO weakens stripes from promoter sites without changing underlying boundaries. (**D)** E-P loop pileup plot using E-P contacts called through H3K27ac and HiC data for 4CBS and Kat6b knockout cells. Red indicates higher contact frequencies on average for E-P loops called, blue indicates lower contact frequency on average. **(E)** P(s) curve showing contact probability at different length scales from HiC data for 4CBS cells (WT) and Kat6b −/− cells. **(F)** Histogram of stripe ratios for Kat6b KO vs control cells for real data compared to randomly generated length ratios. Both distributions are generated using ∼10,000 stripe length pairs. A two-sided t-test was performed, returning a p-value of 1.06e-40. (**G)** ChIP-Seq signal tracks (normalized read counts) of Flag-Kat6b, H3K27ac, CTCF, Rad21 in control and Kat6b −/− cells. (**H)** ChromHMM heatmaps displaying the amount of activity of each histone modification in the identified states, enrichment of genomic regions and Kat6b. **(I)** Flag-Kat6b binding at mouse ESC histone modification peaks (from ENCODE).

To determine the changes to chromatin topology and genome organization upon Kat6b loss, we performed *in situ* Hi-C using Kat6b KO cells (Fig. 6C-F). Contact maps showed less long range looping in Kat6b KO cells (Fig. 6C). Despite the changes in long range looping, insulation scores in Kat6b KO cells compared to 4CBS cells indicated that there is not a significant weakening in insulation strength at TAD boundaries (fig. S7f). When we performed aggregate peak analysis (APA) for 24,353 enhancer-promoter loops, we found a dramatic weakening in contact frequencies in Kat6b KO cells, indicating a likely effect on individual contacts instead of TAD boundaries themselves (Fig. 6D). Confirming the changes in looping, we found that loss of Kat6b resulted in loss of longer range contacts genome-wide (∼100 kb-1 Mb) (Fig. 6E), indicative of a defect in loop extrusion (*49*). To further assess the changes in cohesin loops, we chose to quantify potential changes in its processivity in Kat6b KO and 4CBS cells using Stripenn (*50*) and calculated the distribution of stripe lengths (Fig. 6F). We identified ∼50% fewer stripes in the Kat6b KO (737/1486) and the stripes exhibited a shorter median stripe length (Fig. 6F; fig. S7g). Consistent with this observation, cohesin subunit Rad21 ChIP-Seq using 4CBS control and Kat6b KO cells showed decreased cohesin retention at the Sox2 promoter upon Kat6b loss (Fig. 6G). To further determine how Kat6b counteracts transcriptional insulation, we next investigated the chromatin binding pattern of Kat6b with ChIP-seq experiments using a 4CBS mESC cell line with Kat6b endogenously tagged on the N-terminus with a Flag epitope (fig. S7c). In line with our observation that there was no significant insulation strength weakening at TAD boundaries, we did not observe Flag-Kat6b binding at CTCF peaks (fig. S7h). Interestingly, Kat6b binding was enriched at active or bivalent gene promoters with 20751 of the 30324 Kat6b peaks coinciding with transcriptional start sites (Fig. 6G-I) further supporting its involvement in facilitating enhancer-promoter loops. In short, our results indicated that Kat6b loss impedes cohesin loop extrusion events that normally mediate enhancer-promoter looping to counteract transcriptional insulation.

### Kat6b inhibition re-establishes transcriptional insulation at many genes in Nipbl defective cells

Mutations in insulator sequences and insulator proteins such as Nipbl and CTCF have been observed in developmental disorders and multiple forms of cancer (*51–54*). In particular, insulation defects were suggested as a mechanism of oncogene activation by nearby enhancers in glioma and gastrointestinal stromal tumors (*55*, *56*). We hypothesized that inhibition of Kat6 activity could alleviate some of the insulation defects due to Nipbl loss observed in disease contexts. We used Nipbl loss to model insulation loss phenotypes also observed in Cornelia de Lange syndrome. We then tested whether inhibition of KAT6 HAT activity can modulate insulation defects caused by the loss of cohesin loader Nipbl.

To determine whether KAT6 acetyltransferase activity is needed for its role in antagonizing transcriptional insulation, we used a Kat6 inhibitor (*57*, *58*). We found that treatment with WM-1119, a potent KAT6A/B inhibitor (*57*) specifically enhanced transcriptional insulation at Nipbl depleted cells (Fig. 7A, *top left*). We further confirmed this by knocking out Kat6b in the Nipbl −/− cells (Fig. 7B, *left*). We did not observe any effect on the uninsulated allele, reaffirming that Kat6b activity is only required to overcome the boundary insulation and allow for eGFP expression at the insulator inserted allele. Collectively, these results suggested that Kat6b antagonizes transcriptional insulation through enhancing enhancer-promoter loops across TAD boundaries.

**Figure 7.**
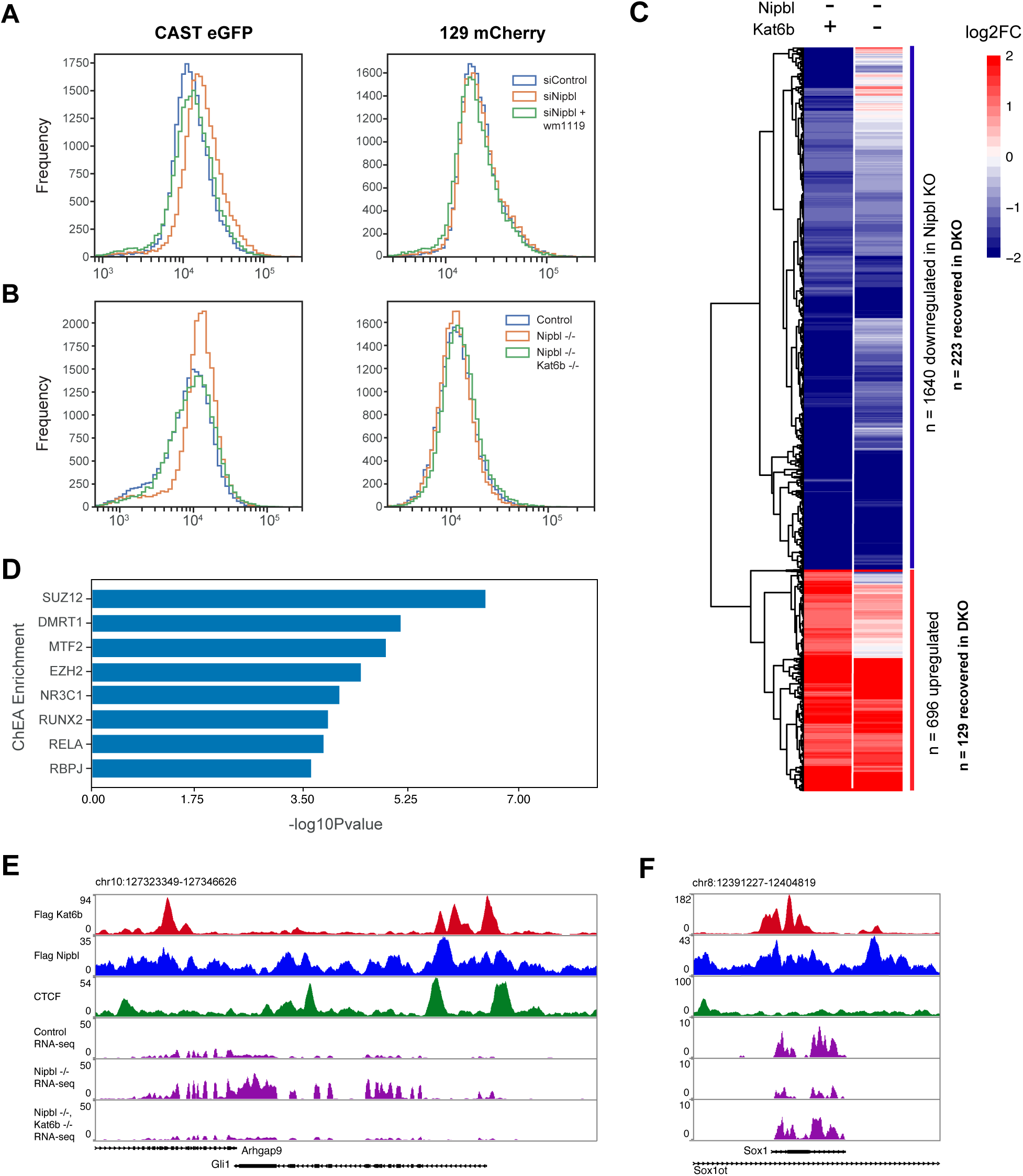
Loss of Kat6b Re-establishes Transcriptional Insulation. **(A)** Flow cytometry histograms for 4CBS cells targeted by siRNAs for control or NipblKO and treated with 10uM WM-1119. **(B)** Flow cytometry histograms for control, Nipbl −/− and Nipbl, Kat6b −/− cells. **(C)** Heatmap of log2 fold changes for a subset DEGs hierarchically clustered based on their fold change over our control for Nipbl −/− and Nipbl−/− combined with Kat6b −/− cell lines. We found n=1640 significantly downregulated genes in Nipbl −/− cells and 693 significantly upregulated genes (l2fc>1, p-value<0.05). Among these genes, 223 saw a transcriptional recovery (no longer significantly changing compared to wild-type) in the downregulated genes, while 129 saw a recovery in the upregulated population. **(D)** ChIP enrichment analysis at rescued genes showing over-representation of factor binding. (**E)** Example Browser track for a gene that is upregulated upon Nipbl loss and rescued upon additional Kat6b −/−. (**F)** Example Browser track for a gene that is downregulated upon Nipbl loss and rescued upon additional Kat6b −/−.

To determine whether Kat6b inhibition could alleviate global insulation defects in Nipbl defective cells, we performed RNA-seq on control, Nipbl −/− and Nipbl, Kat6b −/− ESCs. We identified 696 genes upregulated and 1640 genes downregulated with a log2FC>1 upon Nipbl loss (Fig. 7C). Among these genes, expression of 329 upregulated and 767 downregulated genes was either not differentially expressed in between WT and double knockout cells, or showed a switch in the direction of change (Fig. 7C; fig. S8a, b). The recovered genes showed enrichment for Kat6b binding at their TSS (Fig. 7E, F). Interestingly, the genes that were rescued in Nipbl−/−, Kat6b−/− cells were also enriched for PRC2 complex member Suz12-binding (Fig. 7D). Taken together, the enrichment we observed in chromHMM analyses for Kat6b binding at bivalent gene promoters (Fig. 6D) and the association with PRC2 complex member SUZ12 at recovered genes (Fig. 7D) further supports that Kat6b is involved in regulation of bivalent gene expression in mESCs, in line with recent studies (*59*).

Our results identified the MORF histone acetyltransferase complex functioning as a novel transcriptional insulation regulator that could counteract some of the phenotypic consequences of cohesin loader loss. We further determined that histone modifiers and insulator proteins function in parallel to mediate gene- and context-specific effects of chromatin architecture by identifying an additional insulation antagonizer from our screen, Dot1l (fig. S9a). Inhibition of Dot1l activity has been shown to block abnormal self-renewal due to cohesin loss in acute myeloid leukemia (*60*). In our work, we discovered loss of Dot1l acted similarly, recovering insulation at the Nipbl KO cells, (fig. S9b) and reverting some of the disregulated genes in the Nipbl KO cells (fig. S9c).

## Discussion

In this study, we systematically investigated the role of loop extrusion machinery and chromatin architecture in enhancer-mediated transcriptional activation and transcriptional insulation. We showed a distinction between the steady-state effects of 3D genome disruption (Rad21 or CTCF depletion) and its effects during state transitions, where it displayed a more pronounced effect on transcription genome-wide. In particular, we found that disruption of allele-specific 3D genome conformations in the steady-state did not result in equivalent transcriptional responses in either allele and in fact showed both alleles maintained pre-depletion transcription levels and E-P contact enrichment despite large-scale expansion. Only upon introduction of differentiation media (dynamic transition) did we see transcriptional responses which reflected the disrupted 3D genome structures upon either Rad21 or CTCF depletion.

Through this, we also demonstrated that CTCF and cohesin complex carry out context-specific functions in transcriptional insulation and enhancer-mediated gene activation. We found that 3D genome contexts and their associated responses upon 3D genome disruption are a direct result of a gene’s chromosomal location relative to insulator elements and enhancer elements. Further, we identified that in our 4CBS cells, the 4xCBS insertion acts to create a novel context in the CAST allele which had context-specific responses that were distinct from WT alleles. In particular, we found that our CAST allele was very sensitive to CTCF depletion during dynamic transitions while the 129 allele was not, a direct result of our introduction of the 4xCBS insertion which contained potent CTCF binding sites to disrupt normal E-P contacts. On the other hand, both alleles showed very similar responses to Rad21 depletion due to a dependence on cohesin to mediate E-P contacts and to bring CTCF anchors together. The allelic contexts drive specific patterning in the local histone modifications, compartmentalization, and looping patterns of each allele and in turn tune the response to specific genome disruption events. Our super-resolution chromatin tracing at 5 kb resolution allowed us to detect physical changes to chromatin structure, which are extremely difficult to identify with 3C based methods. Combined with ChIP and gene expression analysis through RNA-seq and flow cytometry, it allowed us to distinguish structure and expression differences upon CTCF removal during different states and across alleles. By utilizing examples of state change such as onset of differentiation and reestablishment of state (A485 recovery), we confirmed that our context dependent transcriptional insulation could not be attributed to changes in the makeup of the culture media alone.

We extrapolated from these two Sox2 contexts to further highlight other unique genes with their own contexts that mirrored or diverged from either Sox2 allele we studied. We highlighted three genes-Nuak2, Islr2, and Trib3. Each has a unique response to the perturbations we performed, with Nuak2 (Fig. 4G) mimicking the Sox2 on the CAST allele due to ectopic activation from a previously less accessible SE, made accessible through weakening of a boundary in the CTCF depletion in differentiation media (boundary located just to the left, outside the window displayed here). Islr2 (fig. S5c) is also at a TAD boundary with multiple CTCF binding sites. These CTCF sites seem to insulate and restrict activation of the SE. Removal of CTCF in both conditions leads to activation of the gene. The removal of Rad21, however, does not affect this, implying additional functions of Rad21 in its regulation. Lastly, Trib3 (fig. S5d), while not affected significantly by CTCF removal, shows a transcriptional reliance on Rad21 in a context-specific manner. In 2i conditions, removal of Rad21 appears to cause a loss of expression. In differentiation media this change is reversed, and loss of Rad21 causes an increase in expression. Within this same genomic region, multiple genes display contextual changes in 2i and differentiation media upon Rad21 depletion.

The context- and gene-specific nature of loop extrusion machinery and chromatin architecture suggested the involvement of additional regulators of transcriptional insulators. To identify such factors, we conducted a genome-wide CRISPR screen using our insulator reporter cells. Through a control screen with cells without 4CBS inserted between Sox2 and the super enhancer, we further distinguished the factors that regulate Sox2 expression in a context and insulation dependent manner. Along with many insulation mediators, we also discovered potential antagonists that weaken transcriptional insulation, including members of the MOZ/ MORF histone acetyltransferase complex (Kat6b, Ing5, Brpf1).

Kat6b and the MORF complex are HATs involved in acetylation of histone lysine residues H3K9, H3K14 and H3K23 and play essential roles in many biological processes including neurogenesis and HSC differentiation (*61*). The significance of Kat6b is underscored by the multiple human developmental disorders caused by Kat6b mutations including Say-Barber-Biesecker-Young variant of Ohdo syndrome (SBBYSS), genitopatellar syndrome (GPS) and Noonan-like syndromes (*62–64*). In addition to human disorders, somatic mutations of the MORF complex have been linked to several cancers including but not limited to breast cancer (*65*), small cell lung cancer (*66*) and AML (*67*). Although recent studies showed promising potential for KAT6 inhibitors in treating various cancers (*58*, *68*), the molecular mechanisms by which Kat6b contributes to these diseases remain an area of active investigation.

In our study, we showed that Kat6b plays a crucial role in facilitating long-range enhancer-promoter loops in mESCs and its loss strengthens transcriptional insulation at the Sox2 locus. We determined that Kat6b binds to active and bivalent TSS - proximal regions. Furthermore, we showed that the acetyltransferase activity of Kat6b is essential for counteracting insulation and that modulation of this activity could be used to neutralize phenotypic consequences of lost insulation, such as those observed in Nipbl deficiency. This observation may have clinical relevance, particularly in developmental cohesinopathies such as Cornelia de Lange Syndrome and certain cancers where the loss of transcriptional insulation can lead to the expression of oncogenes (*55*). Our findings suggest a possible mechanism of action for the recently FDA-approved Kat6 inhibitors in cancer therapy. Corroborating our result, inhibitors for Dot1l, another hit from our screen that binds to bivalent regions (*69*), appears to block self-renewal of cohesin mutant hematopoietic stem and progenitor cells (*60*).

One limitation of our work was that we focused solely on the 4CBS-Sox2 locus in our GECKO screen to identify insulation regulators. Given that CTCF acts in a context and gene specific manner, we suspect using another insulator locus could uncover different or additional regulators. While we discovered that Kat6b antagonizes insulation at the 4CBS insert and facilitates enhancer-promoter looping in mESCs, it is likely that other regulators are involved in different loci or contexts. In agreement with this, an earlier GECKO screen for CTCF insulation regulators at the HoxA locus revealed MAZ and other tissue-specific ZNF proteins (*70*, *71*). Furthermore, another study that employed FISH-based screening to identify TAD boundary regulators among the 3083 druggable genes have highlighted a role for kinase the GSK3A in restricting chromatin looping (*37*). We suspect that one potential reason we did not identify GSK3A as a significant hit in our screen was because our mESC culture media already contained GSK3 inhibitors.

Another limitation of our work was cell cycle specific effects that are seen with long term Rad21 depletion factoring into our genome wide datasets. Shorter depletions of Rad21 resulted in a comparable number of differentially expressed genes to earlier studies (*23*). Regardless, in any Rad21 depleted perturbation, we observed a subtle increase in both eGFP and mCherry expression, even during p300 inhibition. We suspect that this is due to increased compartmentalization and formation of large accessible chromatin domains which allow Sox2 to be in vicinity to other enhancers (*24*, *25*).

In summary, our results elucidate the requirements for insulators and enhancer-promoter contacts in transcriptional gene regulation, particularly highlighting the function of CTCF as stabilizer of transcriptional states by delimiting enhancer influence during transcriptional state changes. It further uncovers regulators of CTCF mediated transcriptional insulation, and highlights Kat6b as a novel insulation antagonizer.

## Acknowledgments

We would like to thank members of the Ren laboratory for comments and discussions. This publication includes data generated at the UC San Diego IGM Genomics Center utilizing an Illumina NovaSeq 6000 that was purchased with funding from a National Institutes of Health SIG grant (#S10 OD026929 to Kristen Jepsen).

## Author contributions

Conceptualization: BR, MT, AJ

Investigation: MT, AJ, BS, HH, YX, QZ, YJ

Formal analysis: AJ, BS, MT, HH, EA

Visualization: AJ, MT, BS

Funding acquisition: BR

Writing - original draft: MT, AJ, BR

Writing - review & editing: All authors contributed

## Data and materials availability

Code used in chromatin tracing and downstream analysis can be found at: https://github.com/ajussila/Tastemel-Jussila_2025

Genomic datasets generated in this study are available at NCBI GEO under accession number GSE288605. We used publicly available datasets from GSE153403 and ENCODE (ENCFF001KER, ENCFF001KDQ and ENCFF008XKX) in Fig. 6.

Imaging Data can be found at 4DN Data portal at: https://data.4dnucleome.org/Tastemel-Jussila_2025

## Supplementary Materials

Materials and Methods Figs. S1 to S9

Tables S1 to S2

---

**Table 1.** 4CBS screen MAGeCK high vs. low summary table. A table showing the gene summary output from MAGeCK, giving p-values, ranks, FDR, and RRA scores for each gene in each of GFP low and GFP high groups.

**Table 2.** Out screen MAGeCK high vs. low summary table. A table showing the gene summary output from MAGeCK, giving p-values, ranks, FDR, and RRA scores for each gene in each of GFP low and GFP high groups.

**Figure S1.**
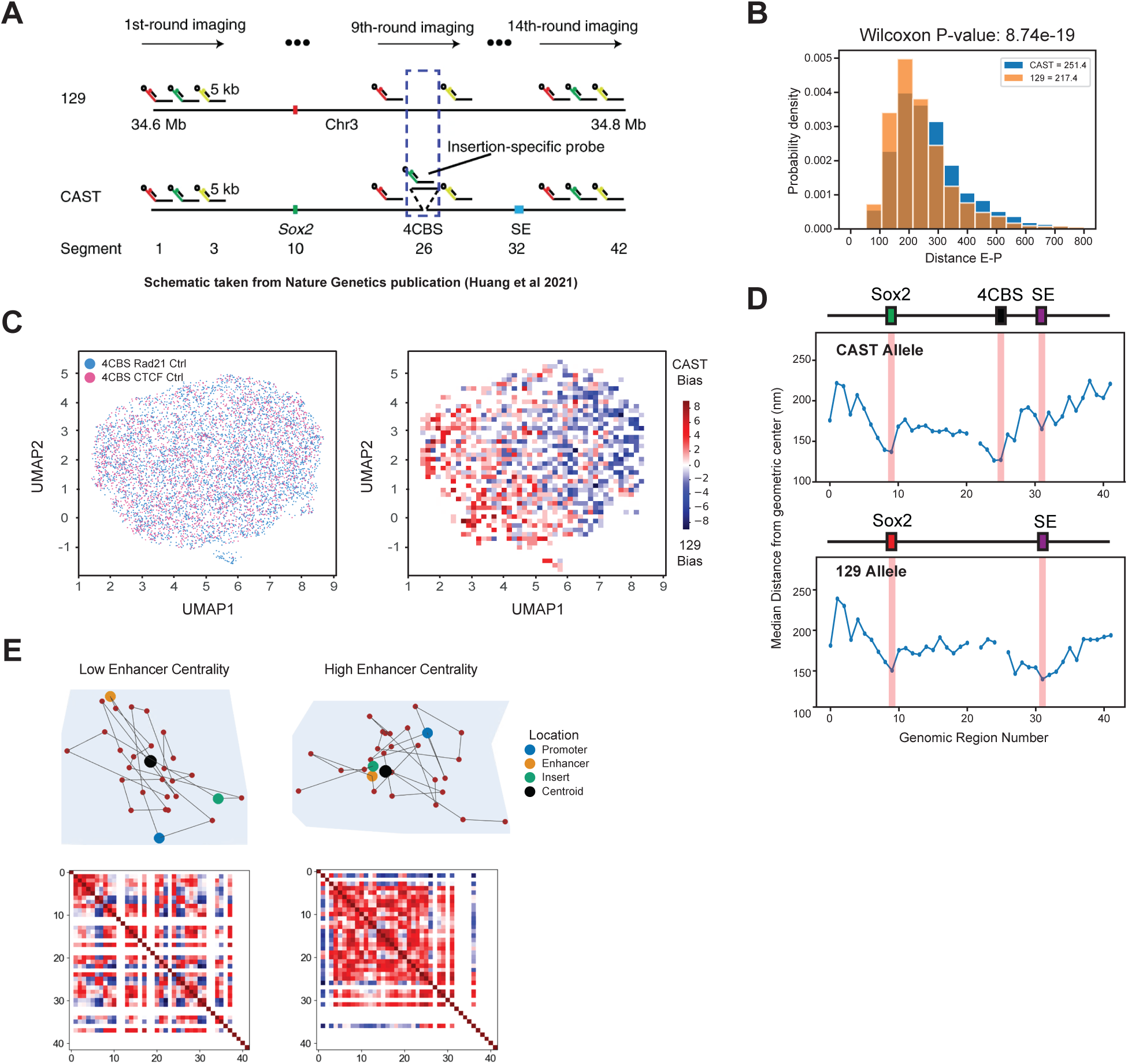
Traditional analyses of chromatin tracing data reveal highly stochastic structures and centrality enrichment of core regulatory elements. **(A)** Schematic depicting probe design scheme for Sox2 locus (212.5 kb on CAST allele and 207.5 kb on 129 allele). Taken directly from our previous publication (*21*). **(B)** E-P distance measurements in single chromosomes of Rad21 degron control cells (no depletion of Rad21). Median distances for CAST and 129 shown in the legend. P-values are computed from Wilcoxon signed rank test. **(C)** UMAP embedding of 4CBS control samples (2 replicates) shows minimal embedding structure (*left*). Downsampled UMAP from previous panel, colored by a ratio of CAST to 129 alleles present in a given bin (*right*). **(D)** Median distance from geometric center across the Sox2 locus, derived from chromatin tracing data. Regions highlighted in red are the promoter (region 9), enhancer (region 31), and insert (region 25). (**E)** Example traces of high and low enhancer centrality chromosomes. *Top*: 3D trace projections showing enhancer, promoter, and insert regions with ball-and-stick style diagrams. *Bottom*: Pairwise distances show high stochasticity in single chromosomes. Missing regions are in white.

**Figure S2.**
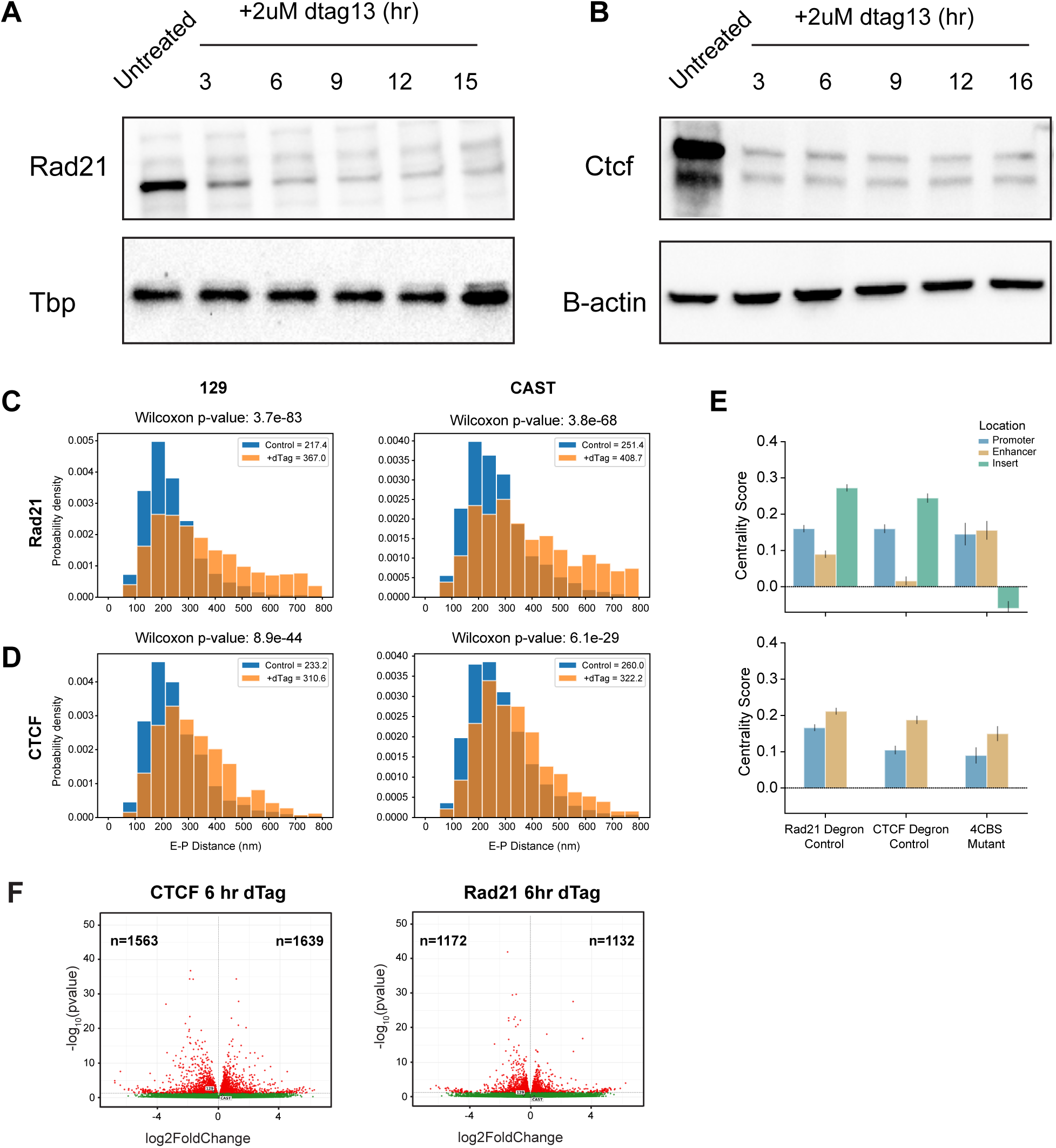
Structural and transcriptional effects of acute CTCF and Rad21 depletion at steady state conditions. **(A)** *Upper:* Western blot results showing Rad21 protein levels prior and after 2uM dTag13 treatments at different time points at Rad21-degron cells. *Lower:* TBP loading control. (**B)** *Upper:* Western blot showing CTCF protein levels before and after addition of 2uM dTag13 to degrade CTCF at various time points. *Lower:* B-Actin loading control. **(C)** Enhancer-promoter distance distributions from chromatin tracing data across alleles and chromosomes for Rad21 control and depletion (+dTag) conditions in 2i media. (**D)** Enhancer-promoter distance distributions from chromatin tracing data across alleles and chromosomes for CTCF control and depletion (+dTag) conditions in 2i media. (**E)** Centrality score measurements showing the Rad21 and CTCF degron control samples compared to the 4CBS mutant (no CTCF binding sites on the 4xCBS insertion) sample to illustrate the conformational change and its dependence on CTCF binding. (**F)** Volcano plots showing the number of significantly changing genes from bulk RNA-seq data in both Rad21 and CTCF 6h depletion conditions. Red dots indicate a significance of p-value < 0.05.

**Figure S3.**
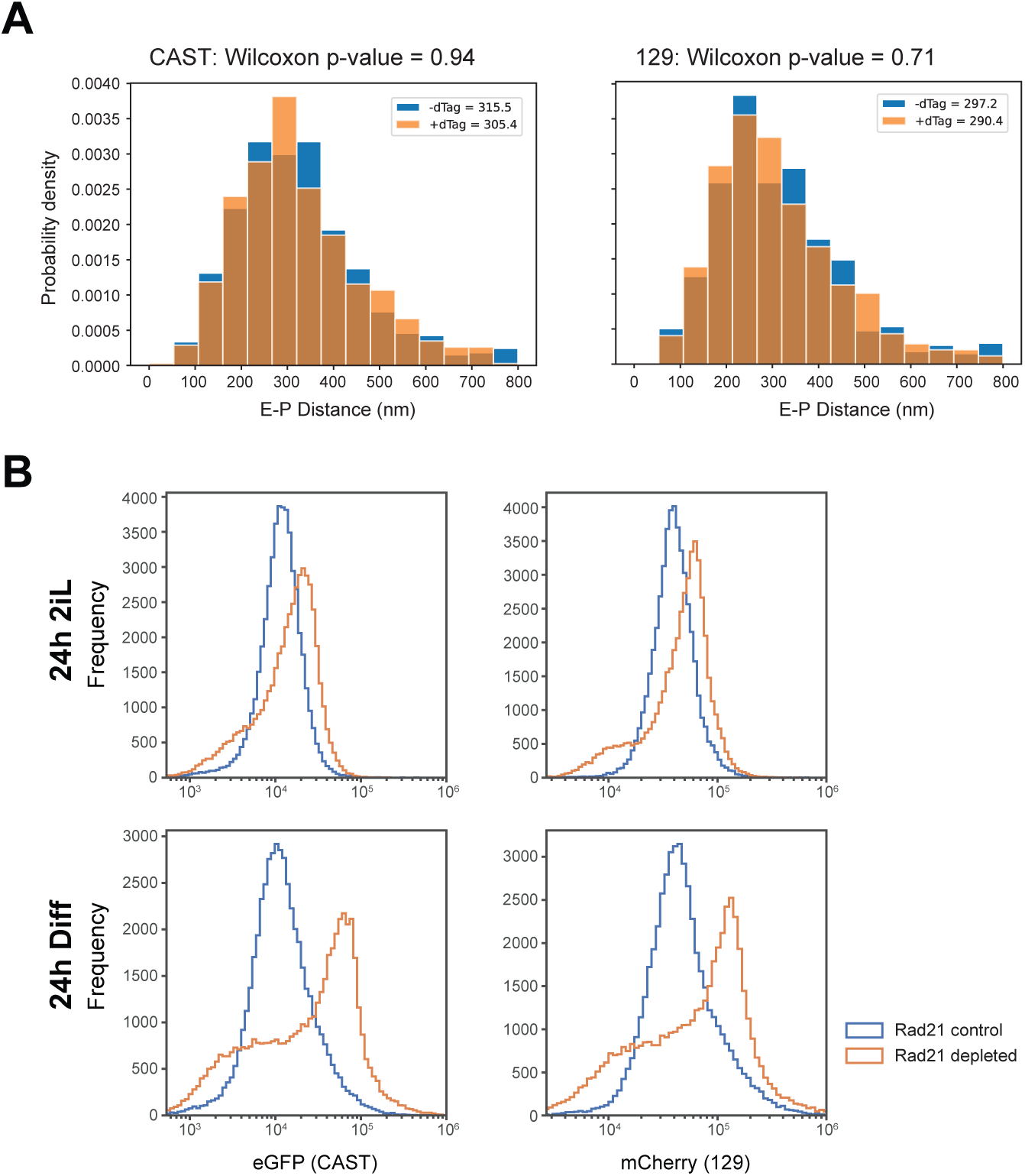
Expression and chromatin structural changes upon onset of differentiation. **(A)** Enhancer-promoter distance histogram from chromatin tracing data showing the distribution of distances across alleles and chromosomes for CTCF control (-dTag) and depletion (+dTag) in differentiation media. (**B)** Histograms of flow cytometry data for 2i and differentiation media (24 hours) with and without dTag in Rad21 degron 4CBS cells.

**Figure S4.**
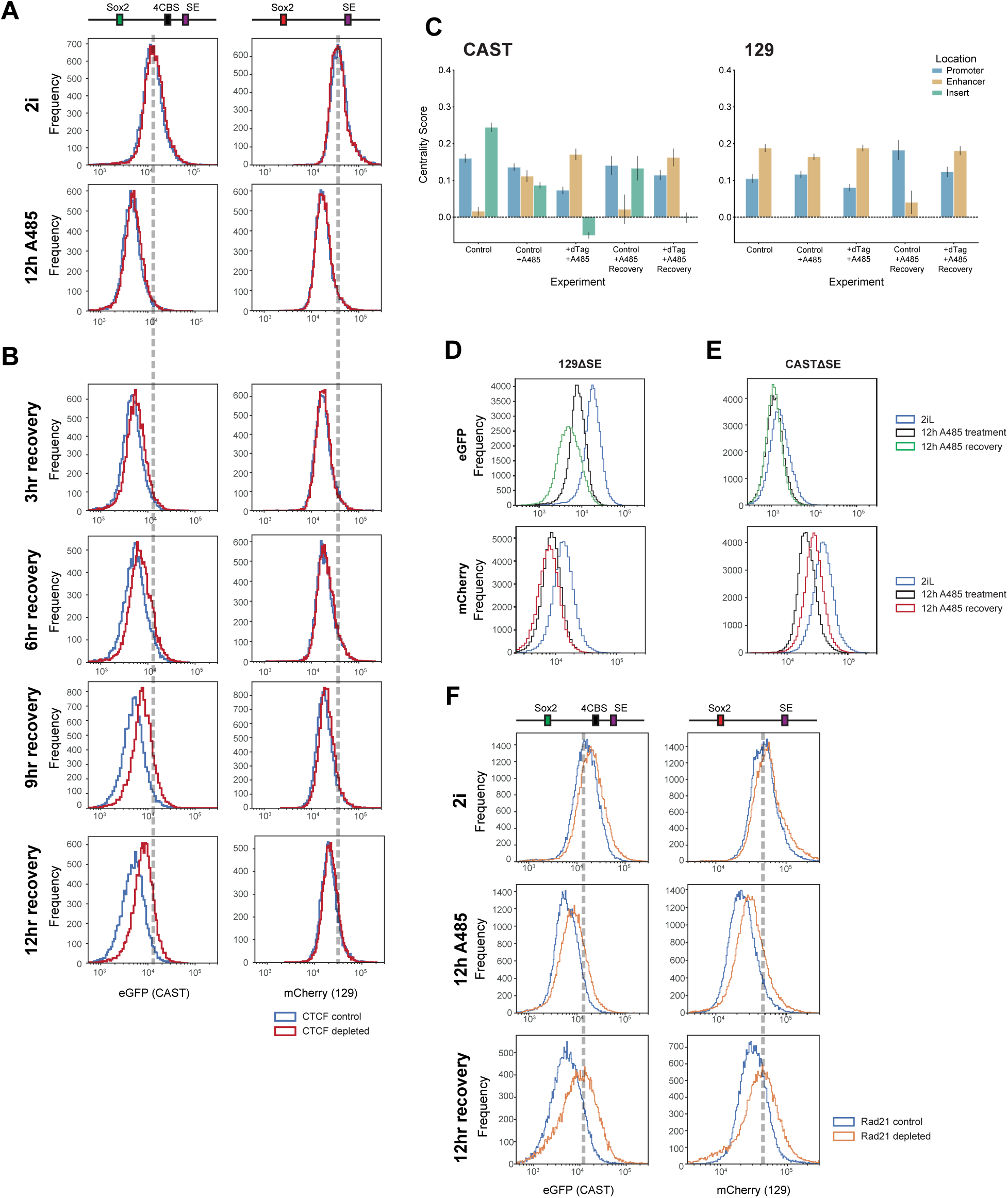
Sox2 expression is reliant on p300 activity at the super enhancer for normal expression. **(A)** Flow cytometry data in 4CBS cells per allele showing WT and A485 treatments. (**B)** Flow cytometry data in 4CBS cells per allele showing recovery time course (3, 6, 9, 12 hrs) after 12 hour A485 treatment to illustrate transcriptional recovery of each allele. **(C)** Bar plot showing median centrality scores for each condition described in panel (3D, E) for enhancer, promoter, and insert regions on each allele. (**D)** Flow cytometry data for 129 allele super-enhancer deleted 4CBS cells in control conditions, 12 hour A485 treatment, and 12 hour A485 treatment plus 6 hour recovery. **(E)** Flow cytometry data for CAST allele super-enhancer deleted 4CBS cells in control conditions, 12 hour A485 treatment, and 12 hour A485 treatment plus 6 hour recovery. **(F)** Flow cytometry data in Rad21 degron 4CBS cells per allele showing control, 12 hour A485 treatment, and 12 hour A485 recovery conditions in the presence and absence of Rad21.

**Figure S5.**
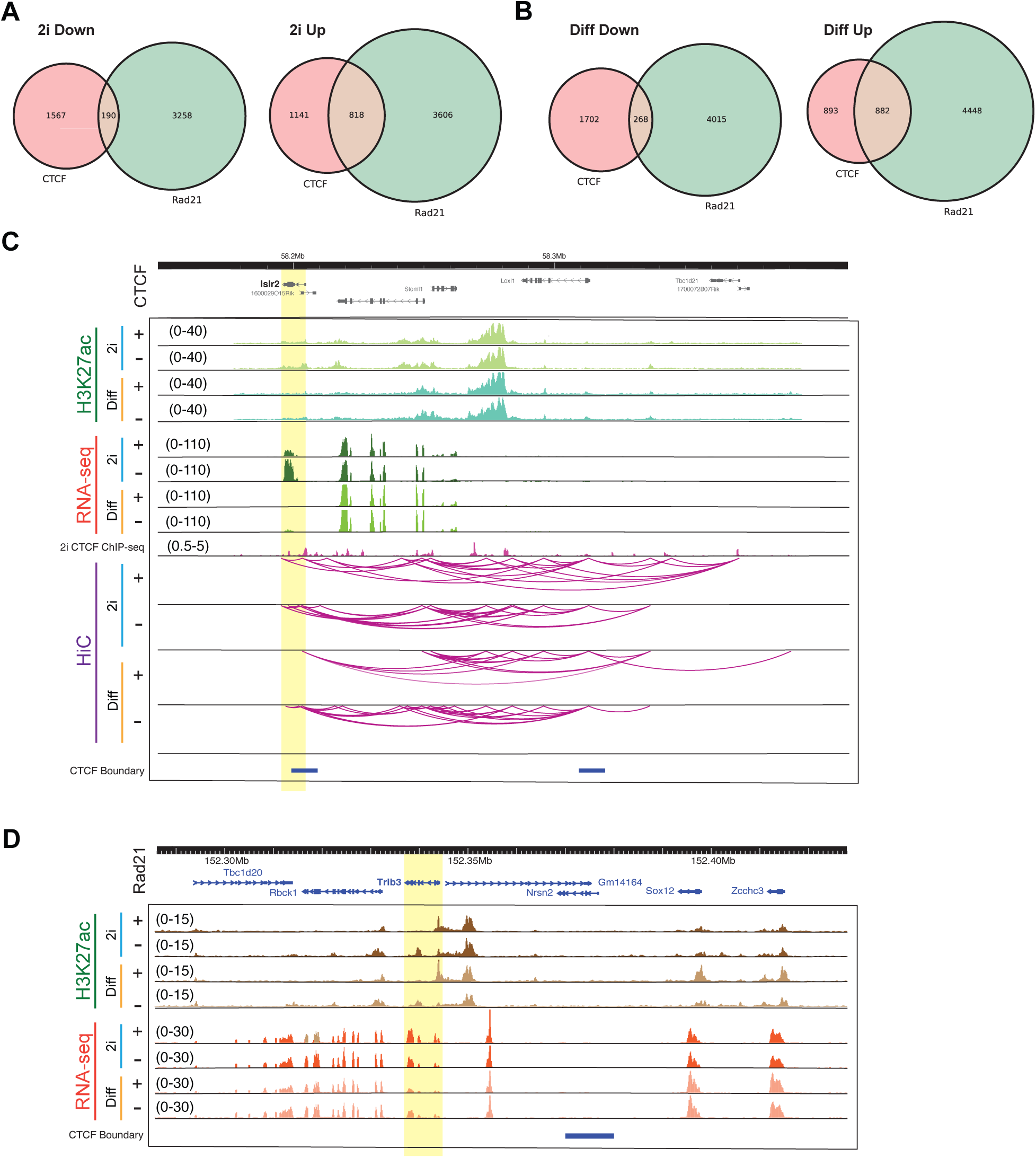
Rad21 and CTCF depletion show overlap primarily in genes induced upon depletion of either factor. **(A)** Venn diagram showing the number of genes which change in the same direction in 2i media between Rad21 and CTCF degron cells. *Left:* Overlaps for downregulated genes in both conditions. *Right:* Overlaps for upregulated genes in both conditions. **(B)** Venn diagram showing the number of genes which change in the same direction in differentiation media between Rad21 and CTCF degron cells. *Left:* Overlaps for genes downregulated in both conditions. *Right:* Overlaps for genes upregulated in both conditions. **(C)** Genome browser tracks showing H3K27ac ChIP-seq, RNA-seq, and inferred HiC promoter contacts for the gene Islr2 with a strong contextual dependence on CTCF. Each set of four tracks are showing 2i, 2i+dTag, differentiation, and differentiation+dTag conditions. In parentheses are the data ranges for the tracks shown. **(D)** Genome browser tracks showing H3K27ac ChIP-seq and RNA-seq for the gene Trib3 with a strong contextual dependence on Rad21. Each set of four tracks are showing 2i, 2i+dTag, differentiation, and differentiation+dTag conditions. Highlighted in yellow is Trib3, which shows a unique contextual dependence on Rad21 for expression. In parentheses are the data ranges for the tracks shown.

**Figure S6.**
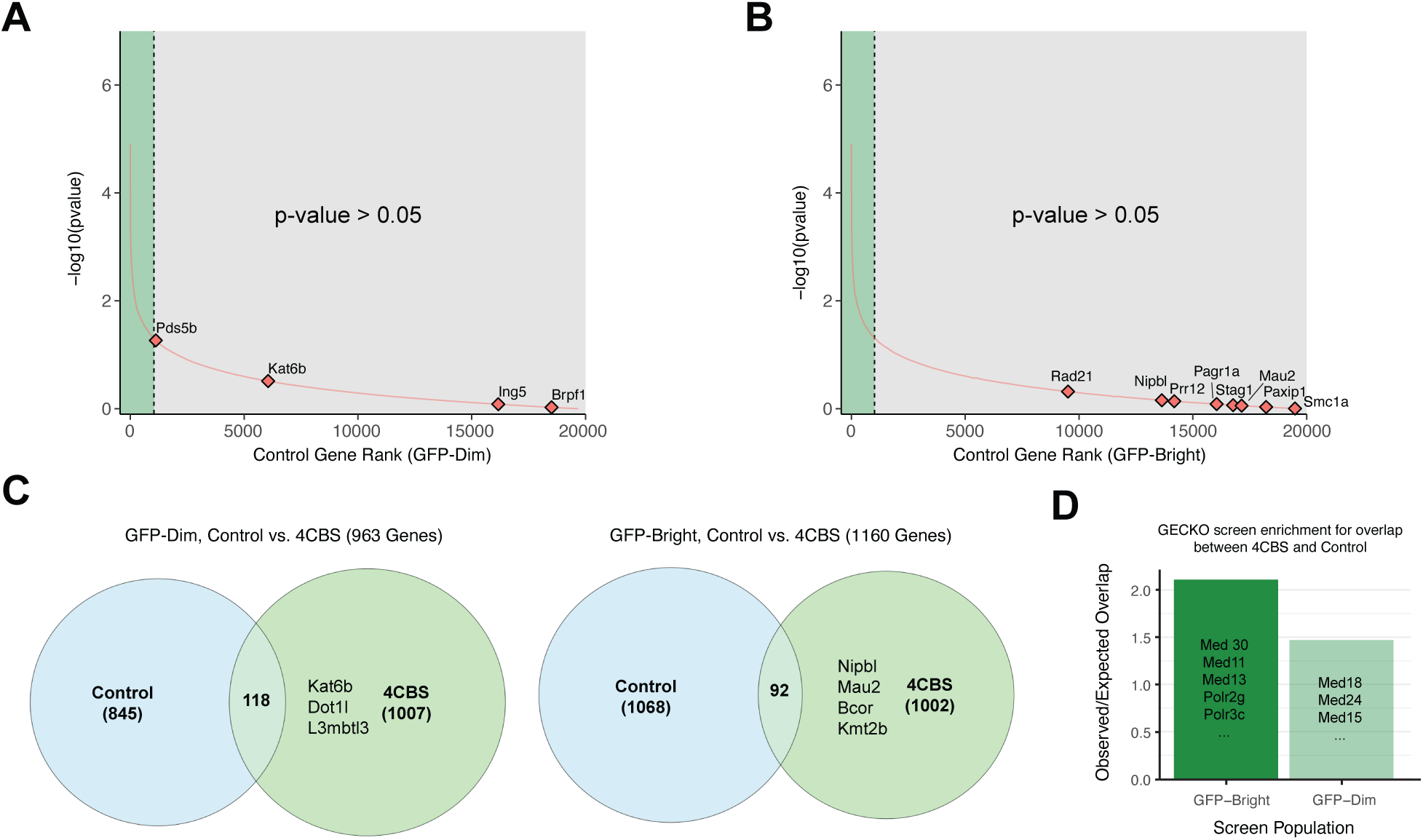
Genome-wide CRISPR Control Screen Distinguished the insulation specific 4CBS screen hits. **(A)** Rank plot of GFP-DIM population (knockout decreases Sox2 expression, weakens insert insulator), showing gene rank vs. p-value for 4CBS-out control cell line, highlighting MORF proteins. Green indicates p-value < 0.05 significance threshold, gray represents non significant hits. (**B)** Rank plot of GFP-BRIGHT population (knockout increases Sox2 expression, strengthens insert insulator), showing gene rank vs. p-value for 4CBS-out control cell line, highlighting insulator proteins. Green indicates p-value < 0.05 significance threshold, gray represents not significant hits. **(C)** Venn diagrams showing overlap between significant genes in 4CBS and 4CBS-out control cell lines in bright and dim populations. Several example genes are listed in the 4CBS-specific group to illustrate which genes are likely insulation-specific in their effects. **(D)** Barplot showing fold enrichment over random expectation (computed through bootstrapping). Enriched gene examples are listed and highlight the transcription-specific roles of many genes shared between 4CBS and 4CBS-out control samples.

**Figure S7.**
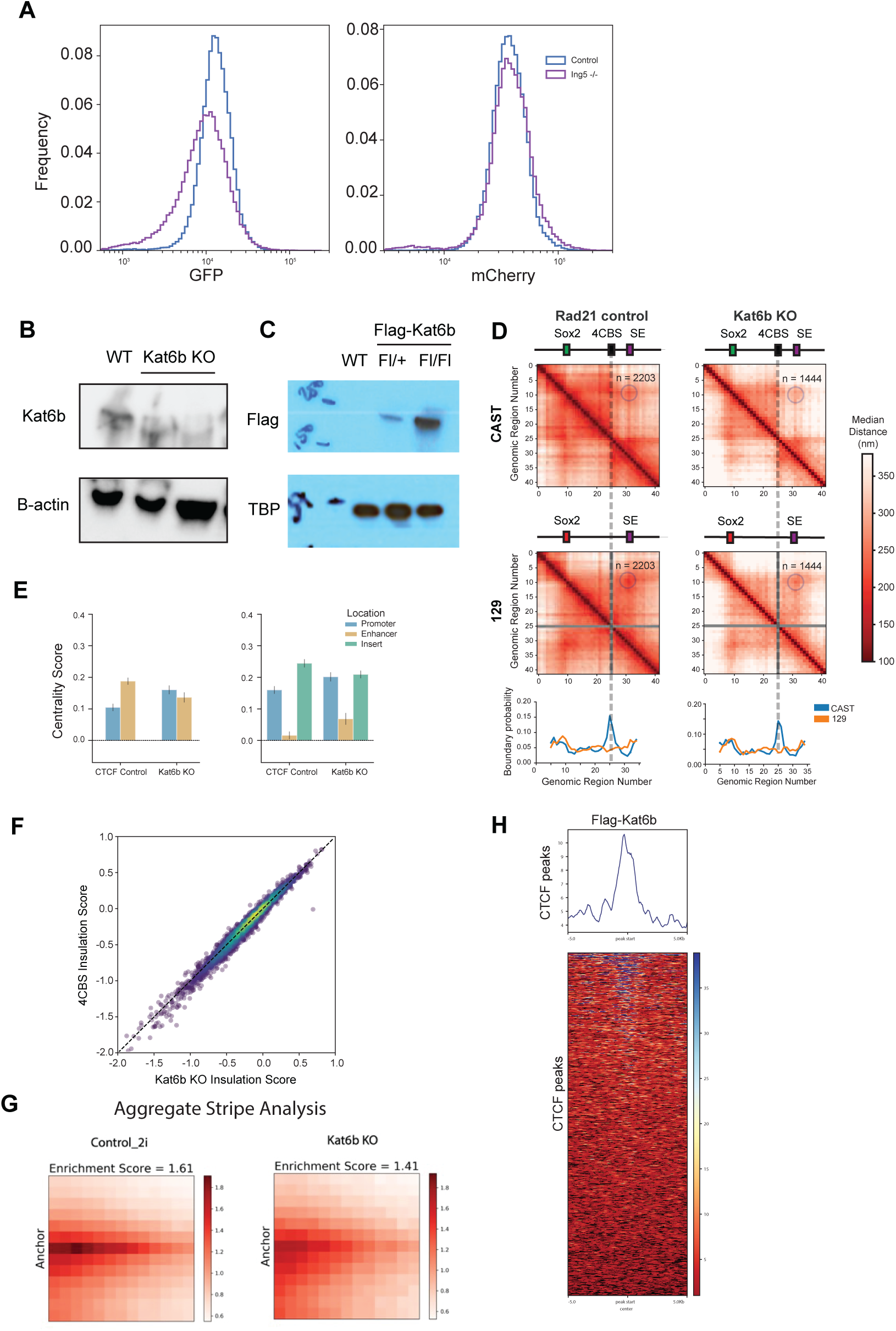
Influence of Kat6b loss on chromatin organization. **(A)** eGFP and mCherry flow cytometry histograms for control and Ing5 −/− 4CBS cells. **(B)** *Top:* Western blot for Kat6b in WT cells and Kat6b knockout cells. *Bottom:* B-actin loading control. **(C)** *Top:* Western blot for Flag in WT cells and heterozygous and homozygous Flag tagged Kat6b cells. *Bottom:* TBP loading control. **(D)** *Upper:* Median distance matrix for CAST and 129 alleles in Rad21 control and Kat6b Knockout cells. *Lower:* Boundary probability computed for each allele from chromatin tracing data. **(E)** Bar plot showing median centrality scores (derived from single chromosome centrality score distributions) per condition for enhancer, promoter, and insert regions on each allele. **(F)** Scatterplot of insulation score calculations derived from HiC data for Kat6b knockout and 4CBS cells using TAD boundaries called in 4CBS cells. The dashed line represents the line x=y to highlight genome-wide changes in boundary strengths between conditions. (**G)** Pileup analysis using Stripenn stripe calls to show changes in stripe length and strength between 4CBS cells and Kat6b knockout cells. Stripe enrichment scores are computed using the Stripenn software. **(H)** Flag-Kat6b binding at mouse ESC CTCF peaks.

**Figure S8.**
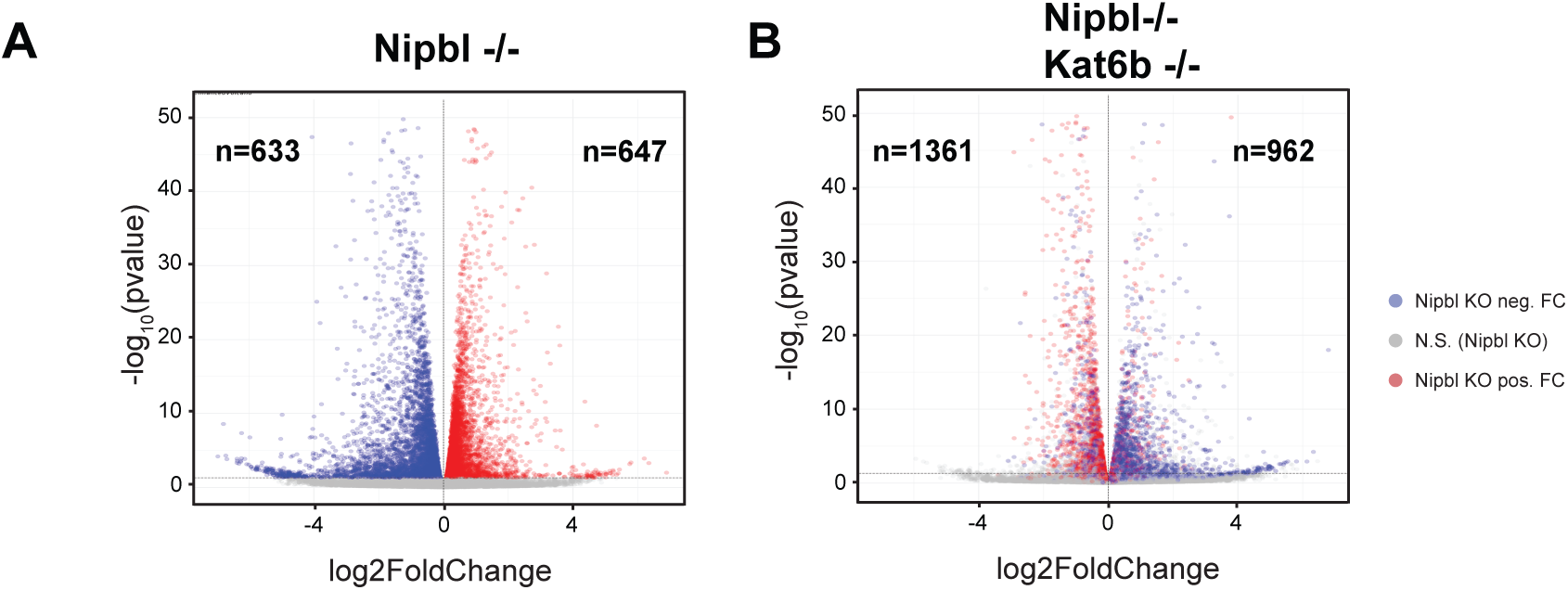
RNA seq analysis of Nipbl −/− and Nipbl,Kat6b double −/− cells. **(A)** Volcano plots showing the number of significantly changing genes from bulk RNA-seq data upon Nipbl −/−. Blue dots indicate genes with p-value < 0.05 and log2FC>1. Genes which were seen to recover upon addition of Kat6b −/− are shown in red. **(B)** Volcano plots showing the number of significantly changing genes from bulk RNA-seq data upon Nipbl −/− and additional Kat6b −/−. Blue dots indicate genes with p-value < 0.05 and log2FC>1. Genes which were seen to recover upon addition of Kat6b −/− are shown in red.

**Figure S9.**
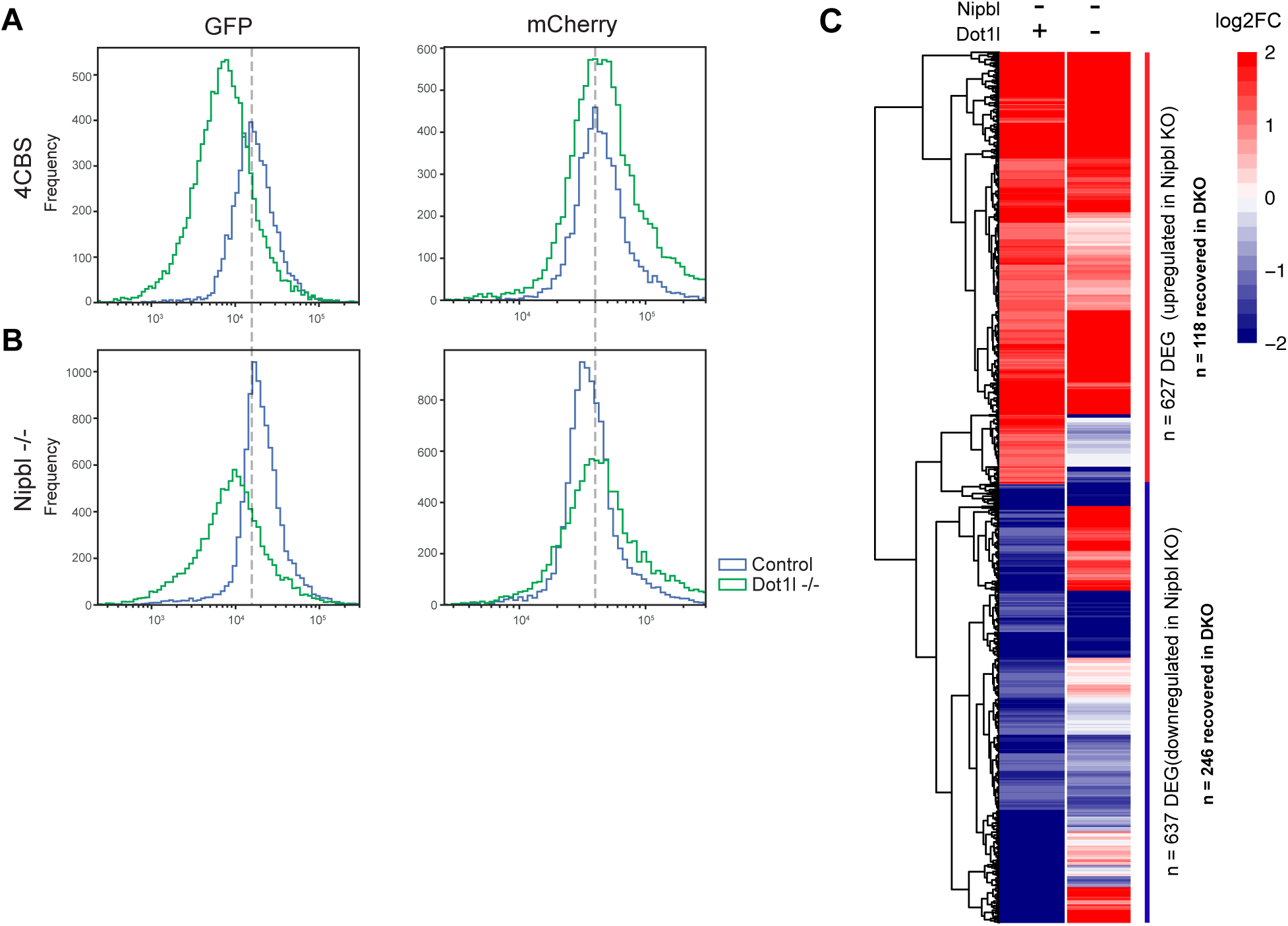
RNA seq analysis of Nipbl −/− and Nipbl,Dot1l double −/− cells. **(A)** Flow cytometry histograms for 4CBS control cells and Dot1l −/− cells for CAST (eGFP) and 129 (mCherry) alleles. **(B)** Flow cytometry histograms for Nipbl −/− cells with and without and Dot1l −/− cells for CAST (eGFP) and 129 (mCherry) alleles. **(C)** Heatmap of log2 fold changes for a subset DEGs hierarchically clustered based on their fold change over our control for Nipbl −/− and Nipbl−/− combined with Dot1l −/− cell lines. We found n=637 significantly downregulated genes in Nipbl −/− cells and 627 significantly upregulated genes (l2fc>1, p-value<0.05). Among these genes, 246 saw a transcriptional recovery in the downregulated genes, while 118 saw a recovery in the upregulated population.

## References and Notes

1. D. Hnisz, D. S. Day, R. A. Young, Insulated neighborhoods: Structural and functional units of mammalian gene control. Cell 167, 1188–1200 (2016).

2. T. Fukaya, B. Lim, M. Levine, Enhancer control of transcriptional bursting. Cell 166, 358–368 (2016).

3. C. R. Bartman, S. C. Hsu, C. C.-S. Hsiung, A. Raj, G. A. Blobel, Enhancer regulation of transcriptional bursting parameters revealed by forced chromatin looping. Mol. Cell 62, 237–247 (2016).

4. W. Deng, J. Lee, H. Wang, J. Miller, A. Reik, P. D. Gregory, A. Dean, G. A. Blobel, Controlling long-range genomic interactions at a native locus by targeted tethering of a looping factor. Cell 149, 1233–1244 (2012).

5. W. Deng, J. W. Rupon, I. Krivega, L. Breda, I. Motta, K. S. Jahn, A. Reik, P. D. Gregory, S. Rivella, A. Dean, G. A. Blobel, Reactivation of developmentally silenced globin genes by forced chromatin looping. Cell 158, 849–860 (2014).

6. B. Tolhuis, R. J. Palstra, E. Splinter, F. Grosveld, W. de Laat, Looping and interaction between hypersensitive sites in the active beta-globin locus. Mol. Cell 10, 1453–1465 (2002).

7. D. Vernimmen, W. A. Bickmore, The hierarchy of transcriptional activation: From enhancer to promoter. Trends Genet. 31, 696–708 (2015).

8. A. L. Sanborn, S. S. P. Rao, S.-C. Huang, N. C. Durand, M. H. Huntley, A. I. Jewett, I. D. Bochkov, D. Chinnappan, A. Cutkosky, J. Li, K. P. Geeting, A. Gnirke, A. Melnikov, D. McKenna, E. K. Stamenova, E. S. Lander, E. L. Aiden, Chromatin extrusion explains key features of loop and domain formation in wild-type and engineered genomes. Proc. Natl. Acad. Sci. U. S. A. 112, E6456–65 (2015).

9. G. Fudenberg, M. Imakaev, C. Lu, A. Goloborodko, N. Abdennur, L. A. Mirny, Formation of chromosomal domains by loop extrusion. Cell Rep. 15, 2038–2049 (2016).

10. S. S. P. Rao, M. H. Huntley, N. C. Durand, E. K. Stamenova, I. D. Bochkov, J. T. Robinson, A. L. Sanborn, I. Machol, A. D. Omer, E. S. Lander, E. L. Aiden, A 3D map of the human genome at kilobase resolution reveals principles of chromatin looping. Cell 159, 1665– 1680 (2014).

11. L. Vian, A. Pękowska, S. S. P. Rao, K.-R. Kieffer-Kwon, S. Jung, L. Baranello, S.-C. Huang, L. El Khattabi, M. Dose, N. Pruett, A. L. Sanborn, A. Canela, Y. Maman, A. Oksanen, W. Resch, X. Li, B. Lee, A. L. Kovalchuk, Z. Tang, S. Nelson, M. Di Pierro, R. R. Cheng, I. Machol, B. G. St Hilaire, N. C. Durand, M. S. Shamim, E. K. Stamenova, J. N. Onuchic, Y. Ruan, A. Nussenzweig, D. Levens, E. L. Aiden, R. Casellas, The energetics and physiological impact of cohesin extrusion. Cell 173, 1165–1178.e20 (2018).

12. W. Schwarzer, N. Abdennur, A. Goloborodko, A. Pekowska, G. Fudenberg, Y. Loe-Mie, N. A. Fonseca, W. Huber, C. H. Haering, L. Mirny, F. Spitz, Two independent modes of chromatin organization revealed by cohesin removal. Nature 551, 51–56 (2017).

13. M. Du, S. H. Stitzinger, J.-H. Spille, W.-K. Cho, C. Lee, M. Hijaz, A. Quintana, I. I. Cissé, Direct observation of a condensate effect on super-enhancer controlled gene bursting. Cell 187, 331–344.e17 (2024).

14. M. Gaszner, G. Felsenfeld, Insulators: exploiting transcriptional and epigenetic mechanisms. Nat. Rev. Genet. 7, 703–713 (2006).

15. T.-H. S. Hsieh, C. Cattoglio, E. Slobodyanyuk, A. S. Hansen, X. Darzacq, R. Tjian, Enhancer–promoter interactions and transcription are largely maintained upon acute loss of CTCF, cohesin, WAPL or YY1. Nat. Genet. 54, 1919–1932 (2022).

16. B. Bintu, L. J. Mateo, J.-H. Su, N. A. Sinnott-Armstrong, M. Parker, S. Kinrot, K. Yamaya, A. N. Boettiger, X. Zhuang, Super-resolution chromatin tracing reveals domains and cooperative interactions in single cells. Science 362 (2018).

17. M. Gabriele, H. B. Brandão, S. Grosse-Holz, A. Jha, G. M. Dailey, C. Cattoglio, T.-H. S. Hsieh, L. Mirny, C. Zechner, A. S. Hansen, Dynamics of CTCF- and cohesin-mediated chromatin looping revealed by live-cell imaging. Science 376, 496–501 (2022).

18. J. Zuin, G. Roth, Y. Zhan, J. Cramard, J. Redolfi, E. Piskadlo, P. Mach, M. Kryzhanovska, G. Tihanyi, H. Kohler, M. Eder, C. Leemans, B. van Steensel, P. Meister, S. Smallwood, L. Giorgetti, Nonlinear control of transcription through enhancer-promoter interactions. Nature 604, 571–577 (2022).

19. A. Hafner, M. Park, S. E. Berger, S. E. Murphy, E. P. Nora, A. N. Boettiger, Loop stacking organizes genome folding from TADs to chromosomes. Mol. Cell 83, 1377–1392.e6 (2023).

20. T.-C. Hung, D. M. Kingsley, A. N. Boettiger, Boundary stacking interactions enable cross-TAD enhancer–promoter communication during limb development. Nat. Genet. 56, 306– 314 (2024).

21. H. Huang, Q. Zhu, A. Jussila, Y. Han, B. Bintu, C. Kern, M. Conte, Y. Zhang, S. Bianco, A. M. Chiariello, M. Yu, R. Hu, M. Tastemel, I. Juric, M. Hu, M. Nicodemi, X. Zhuang, B. Ren, CTCF mediates dosage- and sequence-context-dependent transcriptional insulation by forming local chromatin domains. Nat. Genet. 53, 1064–1074 (2021).

22. E. P. Nora, A. Goloborodko, A.-L. Valton, J. H. Gibcus, A. Uebersohn, N. Abdennur, J. Dekker, L. A. Mirny, B. G. Bruneau, Targeted Degradation of CTCF Decouples Local Insulation of Chromosome Domains from Genomic Compartmentalization. Cell 169, 930–944.e22 (2017).

23. S. S. P. Rao, S.-C. Huang, B. Glenn St Hilaire, J. M. Engreitz, E. M. Perez, K.-R. Kieffer-Kwon, A. L. Sanborn, S. E. Johnstone, G. D. Bascom, I. D. Bochkov, X. Huang, M. S. Shamim, J. Shin, D. Turner, Z. Ye, A. D. Omer, J. T. Robinson, T. Schlick, B. E. Bernstein, R. Casellas, E. S. Lander, E. L. Aiden, Cohesin Loss Eliminates All Loop Domains. Cell 171, 305–320.e24 (2017).

24. L. Xie, P. Dong, Y. Qi, T.-H. S. Hsieh, B. P. English, S. Jung, X. Chen, M. De Marzio, R. Casellas, H. Y. Chang, B. Zhang, R. Tjian, Z. Liu, BRD2 compartmentalizes the accessible genome. Nat. Genet. 54, 481–491 (2022).

25. P. Dong, S. Zhang, V. Gandin, L. Xie, L. Wang, A. L. Lemire, W. Li, H. Otsuna, T. Kawase, A. D. Lander, H. Y. Chang, Z. J. Liu, Cohesin prevents cross-domain gene coactivation. Nat. Genet. 56, 1654–1664 (2024).

26. N. Kubo, H. Ishii, X. Xiong, S. Bianco, F. Meitinger, R. Hu, J. D. Hocker, M. Conte, D. Gorkin, M. Yu, B. Li, J. R. Dixon, M. Hu, M. Nicodemi, H. Zhao, B. Ren, Promoter-proximal CTCF binding promotes distal enhancer-dependent gene activation. Nat. Struct. Mol. Biol. 28, 152–161 (2021).

27. Y. Cheng, M. Liu, M. Hu, S. Wang, TAD-like single-cell domain structures exist on both active and inactive X chromosomes and persist under epigenetic perturbations. Genome Biol. 22, 309 (2021).

28. D. G. Lupiáñez, K. Kraft, V. Heinrich, P. Krawitz, F. Brancati, E. Klopocki, D. Horn, H. Kayserili, J. M. Opitz, R. Laxova, F. Santos-Simarro, B. Gilbert-Dussardier, L. Wittler, M. Borschiwer, S. A. Haas, M. Osterwalder, M. Franke, B. Timmermann, J. Hecht, M. Spielmann, A. Visel, S. Mundlos, Disruptions of topological chromatin domains cause pathogenic rewiring of gene-enhancer interactions. Cell 161, 1012–1025 (2015).

29. M. Franke, D. M. Ibrahim, G. Andrey, W. Schwarzer, V. Heinrich, R. Schöpflin, K. Kraft, R. Kempfer, I. Jerković, W.-L. Chan, M. Spielmann, B. Timmermann, L. Wittler, I. Kurth, P. Cambiaso, O. Zuffardi, G. Houge, L. Lambie, F. Brancati, A. Pombo, M. Vingron, F. Spitz, S. Mundlos, Formation of new chromatin domains determines pathogenicity of genomic duplications. Nature 538, 265–269 (2016).

30. J. Ibn-Salem, S. Köhler, M. I. Love, H.-R. Chung, N. Huang, M. E. Hurles, M. Haendel, N. L. Washington, D. Smedley, C. J. Mungall, S. E. Lewis, C.-E. Ott, S. Bauer, P. N. Schofield, S. Mundlos, M. Spielmann, P. N. Robinson, Deletions of chromosomal regulatory boundaries are associated with congenital disease. Genome Biol. 15, 423 (2014).

31. M. Laugsch, M. Bartusel, R. Rehimi, H. Alirzayeva, A. Karaolidou, G. Crispatzu, P. Zentis, M. Nikolic, T. Bleckwehl, P. Kolovos, W. F. J. van Ijcken, T. Šarić, K. Koehler, P. Frommolt, K. Lachlan, J. Baptista, A. Rada-Iglesias, Modeling the pathological long-range regulatory effects of human structural variation with patient-specific hiPSCs. Cell Stem Cell 24, 736–752.e12 (2019).

32. J. H. Sun, L. Zhou, D. J. Emerson, S. A. Phyo, K. R. Titus, W. Gong, T. G. Gilgenast, J. A. Beagan, B. L. Davidson, F. Tassone, J. E. Phillips-Cremins, Disease-associated short tandem repeats co-localize with chromatin domain boundaries. Cell 175, 224–238.e15 (2018).

33. C. Redin, H. Brand, R. L. Collins, T. Kammin, E. Mitchell, J. C. Hodge, C. Hanscom, V. Pillalamarri, C. M. Seabra, M.-A. Abbott, O. A. Abdul-Rahman, E. Aberg, R. Adley, S. L. Alcaraz-Estrada, F. S. Alkuraya, Y. An, M.-A. Anderson, C. Antolik, K. Anyane-Yeboa, J. F. Atkin, T. Bartell, J. A. Bernstein, E. Beyer, I. Blumenthal, E. M. H. F. Bongers, E. H. Brilstra, C. W. Brown, H. T. Brüggenwirth, B. Callewaert, C. Chiang, K. Corning, H. Cox, E. Cuppen, B. B. Currall, T. Cushing, D. David, M. A. Deardorff, A. Dheedene, M. D’Hooghe, B. B. A. de Vries, D. L. Earl, H. L. Ferguson, H. Fisher, D. R. FitzPatrick, P. Gerrol, D. Giachino, J. T. Glessner, T. Gliem, M. Grady, B. H. Graham, C. Griffis, K. W. Gripp, A. L. Gropman, A. Hanson-Kahn, D. J. Harris, M. A. Hayden, R. Hill, R. Hochstenbach, J. D. Hoffman, R. J. Hopkin, M. W. Hubshman, A. M. Innes, M. Irons, M. Irving, J. C. Jacobsen, S. Janssens, T. Jewett, J. P. Johnson, M. C. Jongmans, S. G. Kahler, D. A. Koolen, J. Korzelius, P. M. Kroisel, Y. Lacassie, W. Lawless, E. Lemyre, K. Leppig, A. V. Levin, H. Li, H. Li, E. C. Liao, C. Lim, E. J. Lose, D. Lucente, M. J. Macera, P. Manavalan, G. Mandrile, C. L. Marcelis, L. Margolin, T. Mason, D. Masser-Frye, M. W. McClellan, C. J. Z. Mendoza, B. Menten, S. Middelkamp, L. R. Mikami, E. Moe, S. Mohammed, T. Mononen, M. E. Mortenson, G. Moya, A. W. Nieuwint, Z. Ordulu, S. Parkash, S. P. Pauker, S. Pereira, D. Perrin, K. Phelan, R. E. P. Aguilar, P. J. Poddighe, G. Pregno, S. Raskin, L. Reis, W. Rhead, D. Rita, I. Renkens, F. Roelens, J. Ruliera, P. Rump, S. L. P. Schilit, R. Shaheen, R. Sparkes, E. Spiegel, B. Stevens, M. R. Stone, J. Tagoe, J. V. Thakuria, B. W. van Bon, J. van de Kamp, I. van Der Burgt, T. van Essen, C. M. van Ravenswaaij-Arts, M. J. van Roosmalen, S. Vergult, C. M. L. Volker-Touw, D. P. Warburton, M. J. Waterman, S. Wiley, A. Wilson, M. de la C. A. Yerena-de Vega, R. T. Zori, B. Levy, H. G. Brunner, N. de Leeuw, W. P. Kloosterman, E. C. Thorland, C. C. Morton, J. F. Gusella, M. E. Talkowski, The genomic landscape of balanced cytogenetic abnormalities associated with human congenital anomalies. Nat. Genet. 49, 36–45 (2017).

34. D. ten Berge, W. Koole, C. Fuerer, M. Fish, E. Eroglu, R. Nusse, Wnt signaling mediates self-organization and axis formation in embryoid bodies. Cell Stem Cell 3, 508–518 (2008).

35. M. Kibschull, Differentiating mouse embryonic stem cells into embryoid bodies in AggreWell plates. Cold Spring Harb. Protoc. 2017, db.prot094169 (2017).

36. L. M. Lasko, C. G. Jakob, R. P. Edalji, W. Qiu, D. Montgomery, E. L. Digiammarino, T. M. Hansen, R. M. Risi, R. Frey, V. Manaves, B. Shaw, M. Algire, P. Hessler, L. T. Lam, T. Uziel, E. Faivre, D. Ferguson, F. G. Buchanan, R. L. Martin, M. Torrent, G. G. Chiang, K. Karukurichi, J. W. Langston, B. T. Weinert, C. Choudhary, P. de Vries, J. H. Van Drie, D. McElligott, E. Kesicki, R. Marmorstein, C. Sun, P. A. Cole, S. H. Rosenberg, M. R. Michaelides, A. Lai, K. D. Bromberg, Discovery of a selective catalytic p300/CBP inhibitor that targets lineage-specific tumours. Nature 550, 128–132 (2017).

37. D. S. Park, S. C. Nguyen, R. Isenhart, P. P. Shah, W. Kim, R. J. Barnett, A. Chandra, J. M. Luppino, J. Harke, M. Wai, P. J. Walsh, R. J. Abdill, R. Yang, Y. Lan, S. Yoon, R. Yunker, M. T. Kanemaki, G. Vahedi, J. E. Phillips-Cremins, R. Jain, E. F. Joyce, High-throughput Oligopaint screen identifies druggable 3D genome regulators. Nature 620, 209–217 (2023).

38. J. G. Doench, N. Fusi, M. Sullender, M. Hegde, E. W. Vaimberg, K. F. Donovan, I. Smith, Z. Tothova, C. Wilen, R. Orchard, H. W. Virgin, J. Listgarten, D. E. Root, Optimized sgRNA design to maximize activity and minimize off-target effects of CRISPR-Cas9. Nat. Biotechnol. 34, 184–191 (2016).

39. W. Li, H. Xu, T. Xiao, L. Cong, M. I. Love, F. Zhang, R. A. Irizarry, J. S. Liu, M. Brown, X. S. Liu, MAGeCK enables robust identification of essential genes from genome-scale CRISPR/Cas9 knockout screens. Genome Biol. 15, 554 (2014).

40. I. Mayayo-Peralta, S. Gregoricchio, K. Schuurman, S. Yavuz, A. Zaalberg, A. Kojic, N. Abbott, B. Geverts, S. Beerthuijzen, J. Siefert, T. M. Severson, M. van Baalen, L. Hoekman, C. Lieftink, M. Altelaar, R. L. Beijersbergen, A. B. Houtsmuller, S. Prekovic, W. Zwart, PAXIP1 and STAG2 converge to maintain 3D genome architecture and facilitate promoter/enhancer contacts to enable stress hormone-dependent transcription. Nucleic Acids Res. 51, 9576–9593 (2023).

41. J. J. M. van Schie, K. de Lint, T. M. Molenaar, M. Moronta Gines, J. A. Balk, M. A. Rooimans, K. Roohollahi, G. M. Pai, L. Borghuis, A. R. Ramadhin, F. Corazza, J. C. Dorsman, K. S. Wendt, R. M. F. Wolthuis, J. de Lange, CRISPR screens in sister chromatid cohesion defective cells reveal PAXIP1-PAGR1 as regulator of chromatin association of cohesin. Nucleic Acids Res. 51, 9594–9609 (2023).

42. J.-S. Kim, X. He, J. Liu, Z. Duan, T. Kim, J. Gerard, B. Kim, M. M. Pillai, W. S. Lane, W. S. Noble, B. Budnik, T. Waldman, Systematic proteomics of endogenous human cohesin reveals an interaction with diverse splicing factors and RNA-binding proteins required for mitotic progression. J. Biol. Chem. 294, 8760–8772 (2019).

43. A. L. Nguyen, E. M. Smith, I. M. Cheeseman, Co-essentiality analysis identifies PRR12 as a cohesin interacting protein and contributor to genomic integrity. Dev. Cell 0 (2024).

44. S. Kueng, B. Hegemann, B. H. Peters, J. J. Lipp, A. Schleiffer, K. Mechtler, J.-M. Peters, Wapl controls the dynamic association of cohesin with chromatin. Cell 127, 955–967 (2006).

45. R. Gandhi, P. J. Gillespie, T. Hirano, Human Wapl is a cohesin-binding protein that promotes sister-chromatid resolution in mitotic prophase. Curr. Biol. 16, 2406–2417 (2006).

46. K.-L. Chan, M. B. Roig, B. Hu, F. Beckouët, J. Metson, K. Nasmyth, Cohesin’s DNA exit gate is distinct from its entrance gate and is regulated by acetylation. Cell 150, 961–974 (2012).

47. B. D. Rowland, M. B. Roig, T. Nishino, A. Kurze, P. Uluocak, A. Mishra, F. Beckouët, P. Underwood, J. Metson, R. Imre, K. Mechtler, V. L. Katis, K. Nasmyth, Building sister chromatid cohesion: smc3 acetylation counteracts an antiestablishment activity. Mol. Cell 33, 763–774 (2009).

48. N. Avvakumov, J. Côté, The MYST family of histone acetyltransferases and their intimate links to cancer. Oncogene 26, 5395–5407 (2007).

49. W. Xi, M. A. Beer, Loop competition and extrusion model predicts CTCF interaction specificity. Nat. Commun. 12, 1046 (2021).

50. S. Yoon, A. Chandra, G. Vahedi, Stripenn detects architectural stripes from chromatin conformation data using computer vision. Nat. Commun. 13, 1602 (2022).

51. B. Adane, G. Alexe, B. K. A. Seong, D. Lu, E. E. Hwang, D. Hnisz, C. A. Lareau, L. Ross, S. Lin, F. S. Dela Cruz, M. Richardson, A. S. Weintraub, S. Wang, A. B. Iniguez, N. V. Dharia, A. S. Conway, A. L. Robichaud, B. Tanenbaum, J. M. Krill-Burger, F. Vazquez, M. Schenone, J. N. Berman, A. L. Kung, S. A. Carr, M. J. Aryee, R. A. Young, B. D. Crompton, K. Stegmaier, STAG2 loss rewires oncogenic and developmental programs to promote metastasis in Ewing sarcoma. Cancer Cell 39, 827–844.e10 (2021).

52. M. Di Nardo, M. M. Pallotta, A. Musio, The multifaceted roles of cohesin in cancer. J. Exp. Clin. Cancer Res. 41, 96 (2022).

53. R. Katainen, K. Dave, E. Pitkänen, K. Palin, T. Kivioja, N. Välimäki, A. E. Gylfe, H. Ristolainen, U. A. Hänninen, T. Cajuso, J. Kondelin, T. Tanskanen, J.-P. Mecklin, H. Järvinen, L. Renkonen-Sinisalo, A. Lepistö, E. Kaasinen, O. Kilpivaara, S. Tuupanen, M. Enge, J. Taipale, L. A. Aaltonen, CTCF/cohesin-binding sites are frequently mutated in cancer. Nat. Genet. 47, 818– 821 (2015).

54. M. De Koninck, A. Losada, Cohesin mutations in cancer. Cold Spring Harb. Perspect. Med. 6, a026476 (2016).

55. W. A. Flavahan, Y. Drier, B. B. Liau, S. M. Gillespie, A. S. Venteicher, A. O. Stemmer-Rachamimov, M. L. Suvà, B. E. Bernstein, Insulator dysfunction and oncogene activation in IDH mutant gliomas. Nature 529, 110–114 (2016).

56. W. A. Flavahan, Y. Drier, S. E. Johnstone, M. L. Hemming, D. R. Tarjan, E. Hegazi, S. J. Shareef, N. M. Javed, C. P. Raut, B. K. Eschle, P. C. Gokhale, J. L. Hornick, E. T. Sicinska, G. D. Demetri, B. E. Bernstein, Altered chromosomal topology drives oncogenic programs in SDH-deficient GISTs. Nature 575, 229–233 (2019).

57. J. B. Baell, D. J. Leaver, S. J. Hermans, G. L. Kelly, M. S. Brennan, N. L. Downer, N. Nguyen, J. Wichmann, H. M. McRae, Y. Yang, B. Cleary, H. R. Lagiakos, S. Mieruszynski, G. Pacini, H. K. Vanyai, M. I. Bergamasco, R. E. May, B. K. Davey, K. J. Morgan, A. J. Sealey, B. Wang, N. Zamudio, S. Wilcox, A. L. Garnham, B. N. Sheikh, B. J. Aubrey, K. Doggett, M. C. Chung, M. de Silva, J. Bentley, P. Pilling, M. Hattarki, O. Dolezal, M. L. Dennis, H. Falk, B. Ren, S. A. Charman, K. L. White, J. Rautela, A. Newbold, E. D. Hawkins, R. W. Johnstone, N. D. Huntington, T. S. Peat, J. K. Heath, A. Strasser, M. W. Parker, G. K. Smyth, I. P. Street, B. J. Monahan, A. K. Voss, T. Thomas, Inhibitors of histone acetyltransferases KAT6A/B induce senescence and arrest tumour growth. Nature 560, 253–257 (2018).

58. S. Sharma, C.-Y. Chung, S. Uryu, J. Petrovic, J. Cao, A. Rickard, N. Nady, S. Greasley, E. Johnson, O. Brodsky, S. Khan, H. Wang, Z. Wang, Y. Zhang, K. Tsaparikos, L. Chen, A. Mazurek, J. Lapek, P.-P. Kung, S. Sutton, P. F. Richardson, E. C. Greenwald, S. Yamazaki, R. Jones, K. A. Maegley, P. Bingham, H. Lam, A. E. Stupple, A. Kamal, A. Chueh, A. Cuzzupe, B. J. Morrow, B. Ren, C. Carrasco-Pozo, C. W. Tan, D. D. Bhuva, E. Allan, E. Surgenor, F. Vaillant, H. Pehlivanoglu, H. Falk, J. R. Whittle, J. Newman, J. Cursons, J. P. Doherty, K. L. White, L. MacPherson, M. Devlin, M. L. Dennis, M. K. Hattarki, M. De Silva, M. A. Camerino, M. S. Butler, O. Dolezal, P. Pilling, R. Foitzik, P. A. Stupple, H. R. Lagiakos, S. R. Walker, S. Hediyeh-Zadeh, S. Nuttall, S. K. Spall, S. A. Charman, T. Connor, T. S. Peat, V. M. Avery, Y. E. Bozikis, Y. Yang, M. Zhang, B. J. Monahan, A. K. Voss, T. Thomas, I. P. Street, S.-J. Dawson, M. A. Dawson, G. J. Lindeman, M. J. Davis, J. E. Visvader, T. A. Paul, Discovery of a highly potent, selective, orally bioavailable inhibitor of KAT6A/B histone acetyltransferases with efficacy against KAT6A-high ER+ breast cancer. Cell Chem. Biol. 30, 1191–1210.e20 (2023).

59. E. Bryan, D. Valsakumar, N. J. Idigo, M. Warburton, K. M. Webb, K. A. McLaughlin, C. Spanos, S. Lenci, V. Major, C. Ambrosi, S. Andrews, T. Baubec, J. Rappsilber, P. Voigt, Nucleosomal asymmetry shapes histone mark binding and promotes poising at bivalent domains. Mol. Cell, doi: 10.1016/j.molcel.2024.12.002 (2024).

60. K. E. Heimbruch, J. B. Fisher, C. T. Stelloh, E. Phillips, M. H. Reimer Jr, A. J. Wargolet, A. E. Meyer, K. Pulakanti, A. D. Viny, J. J. Loppnow, R. L. Levine, J. A. Pulikkan, N. Zhu, S. Rao, DOT1L inhibitors block abnormal self-renewal induced by cohesin loss. Sci. Rep. 11, 7288 (2021).

61. G. Zu, Y. Liu, J. Cao, B. Zhao, H. Zhang, L. You, BRPF1-KAT6A/KAT6B complex: Molecular structure, biological function and human disease. Cancers (Basel) 14 (2022).

62. J. Clayton-Smith, J. O’Sullivan, S. Daly, S. Bhaskar, R. Day, B. Anderson, A. K. Voss, T. Thomas, L. G. Biesecker, P. Smith, A. Fryer, K. E. Chandler, B. Kerr, M. Tassabehji, S.-A. Lynch, M. Krajewska-Walasek, S. McKee, J. Smith, E. Sweeney, S. Mansour, S. Mohammed, D. Donnai, G. Black, Whole-exome-sequencing identifies mutations in histone acetyltransferase gene KAT6B in individuals with the Say-Barber-Biesecker variant of Ohdo syndrome. Am. J. Hum. Genet. 89, 675–681 (2011).

63. M. A. Simpson, C. Deshpande, D. Dafou, L. E. L. M. Vissers, W. J. Woollard, S. E. Holder, G. Gillessen-Kaesbach, R. Derks, S. M. White, R. Cohen-Snuijf, S. G. Kant, L. H. Hoefsloot, W. Reardon, H. G. Brunner, E. M. H. F. Bongers, R. C. Trembath, De novo mutations of the gene encoding the histone acetyltransferase KAT6B cause Genitopatellar syndrome. Am. J. Hum. Genet. 90, 290–294 (2012).

64. M. Kraft, I. C. Cirstea, A. K. Voss, T. Thomas, I. Goehring, B. N. Sheikh, L. Gordon, H. Scott, G. K. Smyth, M. R. Ahmadian, U. Trautmann, M. Zenker, M. Tartaglia, A. Ekici, A. Reis, H.-G. Dörr, A. Rauch, C. T. Thiel, Disruption of the histone acetyltransferase MYST4 leads to a Noonan syndrome-like phenotype and hyperactivated MAPK signaling in humans and mice. J. Clin. Invest. 121, 3479–3491 (2011).

65. H. Lynch, H. Wen, Y. C. Kim, C. Snyder, Y. Kinarsky, P. X. Chen, F. Xiao, D. Goldgar, K. H. Cowan, S. M. Wang, Can unknown predisposition in familial breast cancer be family-specific? Breast J. 19, 520–528 (2013).

66. L. Simó-Riudalbas, M. Pérez-Salvia, F. Setien, A. Villanueva, C. Moutinho, A. Martínez-Cardús, S. Moran, M. Berdasco, A. Gomez, E. Vidal, M. Soler, H. Heyn, A. Vaquero, C. de la Torre, S. Barceló-Batllori, A. Vidal, L. Roz, U. Pastorino, K. Szakszon, G. Borck, C. S. Moura, F. Carneiro, I. Zondervan, S. Savola, R. Iwakawa, T. Kohno, J. Yokota, M. Esteller, KAT6B is a tumor suppressor histone H3 lysine 23 acetyltransferase undergoing genomic loss in small cell lung cancer. Cancer Res. 75, 3936–3945 (2015).

67. I. Panagopoulos, T. Fioretos, M. Isaksson, U. Samuelsson, R. Billström, B. Strömbeck, F. Mitelman, B. Johansson, Fusion of the MORF and CBP genes in acute myeloid leukemia with the t(10;16)(q22;p13). Hum. Mol. Genet. 10, 395–404 (2001).

68. T. Mukohara, Y. H. Park, D. Sommerhalder, K. Yonemori, E. Hamilton, S.-B. Kim, J. H. Kim, H. Iwata, T. Yamashita, R. M. Layman, M. Mita, T. Clay, Y. S. Chae, C. Oakman, F. Yan, G. M. Kim, S.-A. Im, G. J. Lindeman, H. S. Rugo, M. Liyanage, M. Saul, C. Le Corre, A. Skoura, L. Liu, M. Li, P. M. LoRusso, Inhibition of lysine acetyltransferase KAT6 in ER+HER2-metastatic breast cancer: a phase 1 trial. Nat. Med. 30, 2242–2250 (2024).

69. B. Appiah, C. L. Fullio, C. Ossola, I. Bertani, E. Restelli, A. Cheffer, M. Polenghi, C. Haffner, M. Garcia-Miralles, P. Zeis, M. Treppner, P. Bovio, L. Schlichtholz, A. Mas-Sanchez, L. Zografidou, J. Winter, H. Binder, D. Grün, N. Kalebic, E. Taverna, T. Vogel, DOT1L activity affects neural stem cell division mode and reduces differentiation and ASNS expression. EMBO Rep. 24, e56233 (2023).

70. H. Ortabozkoyun, P.-Y. Huang, H. Cho, V. Narendra, G. LeRoy, E. Gonzalez-Buendia, J. A. Skok, A. Tsirigos, E. O. Mazzoni, D. Reinberg, CRISPR and biochemical screens identify MAZ as a cofactor in CTCF-mediated insulation at Hox clusters. Nat. Genet. 54, 202–212 (2022).

71. H. Ortabozkoyun, P.-Y. Huang, E. Gonzalez-Buendia, H. Cho, S. Y. Kim, A. Tsirigos, E. O. Mazzoni, D. Reinberg, Members of an array of zinc-finger proteins specify distinct Hox chromatin boundaries. Mol. Cell 84, 3406–3422.e6 (2024).

72. Y. Li, C. M. Rivera, H. Ishii, F. Jin, S. Selvaraj, A. Y. Lee, J. R. Dixon, B. Ren, CRISPR reveals a distal super-enhancer required for Sox2 expression in mouse embryonic stem cells. PLoS One 9, e114485 (2014).

73. B. Teague, Cytoflow: A Python Toolbox for Flow Cytometry, bioRxiv (2022) p. 2022.07.22.501078.

74. K. Zhang, N. R. Zemke, E. J. Armand, B. Ren, A fast, scalable and versatile tool for analysis of single-cell omics data. Nat. Methods 21, 217–227 (2024).

